# Identification and targeting of microbial putrescine acetylation in bloodstream infections

**DOI:** 10.1101/2023.09.21.558834

**Authors:** Jared R. Mayers, Jack Varon, Ruixuan R. Zhou, Martin Daniel-Ivad, Courtney Beaulieu, Amrisha Bholse, Nathaniel R. Glasser, Franziska M. Lichtenauer, Julie Ng, Mayra Pinilla Vera, Curtis Huttenhower, Mark A. Perrella, Clary B. Clish, Sihai D. Zhao, Rebecca M. Baron, Emily P. Balskus

## Abstract

The growth of antimicrobial resistance (AMR) has highlighted an urgent need to identify bacterial pathogenic functions that may be targets for clinical intervention. Although severe bacterial infections profoundly alter host metabolism, prior studies have largely ignored alterations in microbial metabolism in this context. Performing metabolomics on patient and mouse plasma samples, we identify elevated levels of bacterially-derived *N*- acetylputrescine during gram-negative bloodstream infections (BSI), with higher levels associated with worse clinical outcomes. We discover that SpeG is the bacterial enzyme responsible for acetylating putrescine and show that blocking its activity reduces bacterial proliferation and slows pathogenesis. Reduction of SpeG activity enhances bacterial membrane permeability and results in increased intracellular accumulation of antibiotics, allowing us to overcome AMR of clinical isolates both in culture and *in vivo.* This study highlights how studying pathogen metabolism in the natural context of infection can reveal new therapeutic strategies for addressing challenging infections.

## INTRODUCTION

Antimicrobial-resistant (AMR) bacterial infections represent an escalating global public health crisis directly responsible for 1.27 million deaths in 2019^1^, a number predicted to rise to 10 million by 2050^2^. Widespread antibiotic administration during the COVID-19 pandemic only accelerated this trend by selecting for the development of resistance in bacterial infections^3-6^. US deaths from AMR infections rose 15% between 2019 to 2020 alone with 40% of these infections acquired in the hospital^7^.

Bloodstream infection (BSI) is a top cause of mortality attributable to AMR^1^. It is present in the majority of patients with septic shock^8-10^, the most severe form of sepsis associated with low blood pressure and organ damage in response to infection. Septic shock has a morality rate over 40%^11,12^. BSIs are most commonly caused by gram-negative bacteria such as *Escherichia coli*, *Klebsiella pneumoniae,* and *Pseudomonas aeruginosa*^13^. These microbes are members of the World Health Organization Priority 1 group of pathogens, recognized as primary drivers of AMR-associated mortality^1^ and priority targets for antibiotic development. Treatment options for BSI and AMR-infections more broadly remain limited both by a narrow repertoire of targets and challenges in drug development that have slowed the pipeline of new antimicrobial agents^14-16^.

Exploiting disease-associated metabolism for clinical benefit has proven promising in contexts such as cancer^17^ and Mendelian disorders^18^, but this paradigm is underutilized in bacterial infections. While prior studies in the context of BSI have focused on how severe bacterial infections alter host metabolism predominantly as a marker of clinical outcomes^19-23^, these studies have largely ignored microbial metabolic activities. Identifying alterations in bacterial metabolism specifically in the context of infection could highlight previously underappreciated processes that play causal roles in disease and are thus more likely to be targets of effective therapies.

In this study, we analyze samples from gram-negative BSIs in patients and animal models to characterize microbial metabolism in the natural context of infection, hypothesizing this would prioritize microbial metabolic processes involved in pathogenesis for further investigation and define new targets for efforts to combat AMR-associated infections. Using an iterative, comparative metabolomics pipeline, we identify acetylation of putrescine as a prominent microbial metabolic activity in BSIs. Characterizing and inhibiting the bacterial acetyltransferase responsible for this activity revealed a direct impact of this metabolism on bacterial fitness in culture and *in vivo*. Finally, this discovery demonstrated that treatment with an acetyltransferase inhibitor could resensitize multi-drug resistant (MDR) bacteria to existing clinical antibiotics. Overall, we have defined a generalizable workflow for identifying, investigating, and exploiting metabolic vulnerabilities in bacterial pathogens.

## RESULTS

### Comparative metabolomics highlights *N*-acetylputrescine as a probable bacterial-derived metabolite in bloodstream infections

To identify bacterial metabolites elevated in human plasma during infection, we first examined an existing cohort of patient plasma samples and identified a small subset from patients admitted to the intensive care unit (ICU) with culture positive gram-negative BSIs (*Escherichia coli, Klebsiella* spp.*, Pseudomonas* spp.) who had banked blood samples drawn contemporaneously or near-contemporaneously with their positive blood cultures as well from controls admitted to the ICU for other reasons (**Figure S1A-B** and **Table S1**)^23^. Targeted liquid chromatography–mass spectrometry (LC–MS)-based metabolomics revealed significant differences in 62 metabolites (with an FDR<0.05) between the two groups (**Figure 1A** and **Table S2**). To prioritize metabolites for further investigation, we determined the extent to which these 62 metabolites were correlated with clinical outcomes as measured by APACHE II scores^24,25^ and found that elevated levels of eight metabolites were associated with increased mortality (**Figure 1A-B** and **Figure S1C**). Of these eight, elevations in *N*- acetylputrescine were the most significantly different between cases and controls (**Figure 1A**). Both *N*- acetylputrescine and a second elevated metabolite, 4-acetamidobutanoate, arise from acetylation of the polyamine putrescine (**Figure 1A-B**)^26,27^. We did not observe elevations in another mono-acetylated polyamine, *N1*-acetylspermidine. We viewed this unexpected increase in levels of metabolites from a specific pathway, putrescine metabolism (**Figure 1C**) as potentially indicating an important role for this pathway during BSI and prioritized these metabolites for further investigation. Incidentally, our cohort reproduced previously reported significant decreases of multiple lysophosphatidyl choline species in sepsis (**Figure S1D**)^28,29^, although these alterations were not associated with clinical outcomes.

**Figure 1.**
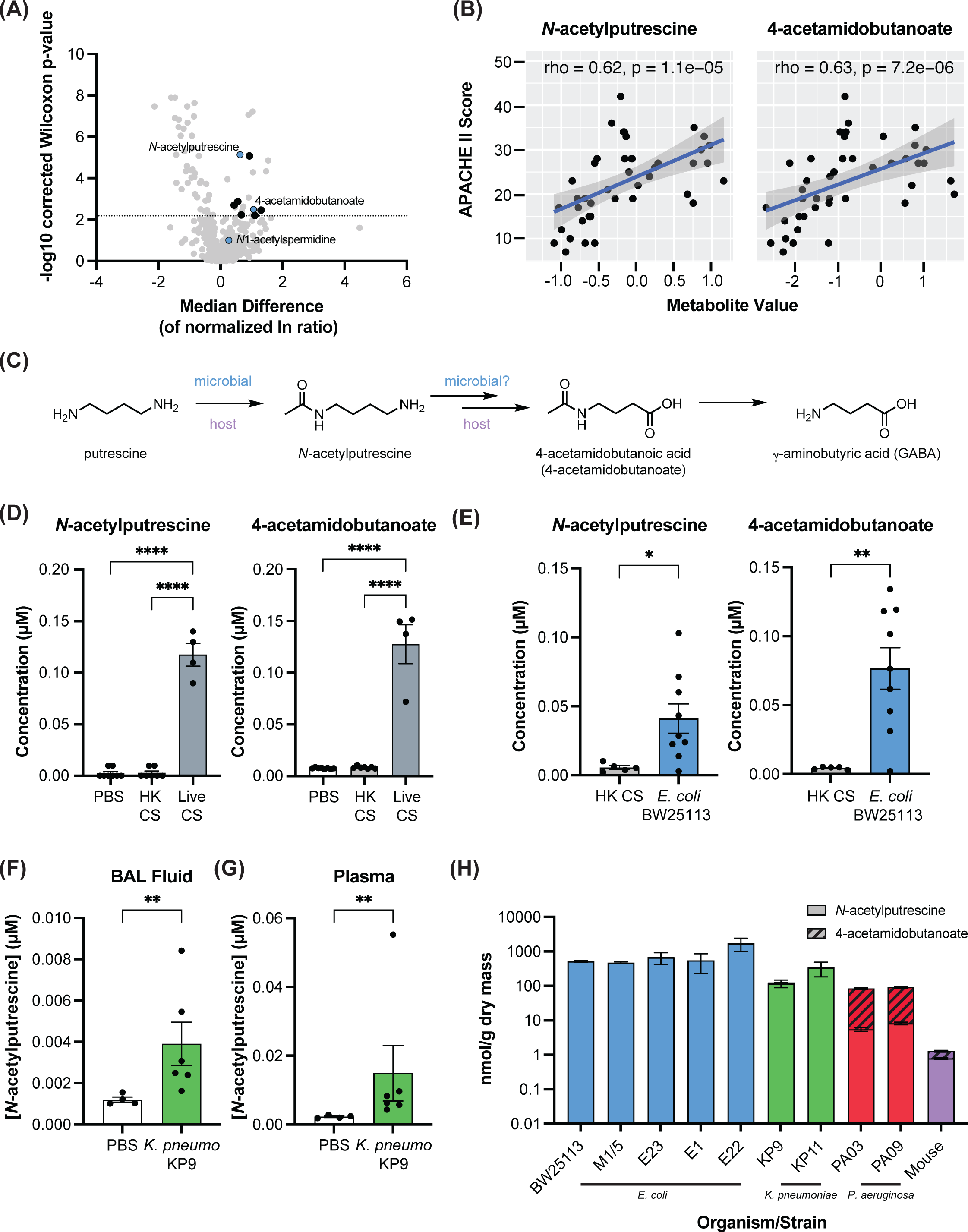
Putrescine metabolites are elevated in humans with gram-negative septic shock and mouse models of sepsis and are produced by bacteria. (A) Volcano plot highlighting significant elevations in *N*-acetylputrescine and 4-acetamidobutanoate in humans with gram-negative septic shock (B) *N*-acetylpturescine and 4-acetomidobutanoate levels correlate with worse clinical outcomes as measured by APACHE II scores (C) Putrescine metabolic pathway outlining production of *N*-acetylputrescine and 4-acetamidobutanoate (D) Putrescine metabolites are elevated in the mouse cecal slurry model of septic shock/BSI; HK = heat killed, CS = cecal slurry; n = 8 PBS, n = 7 HK CS, n = 4 Live CS; p-values were determined by One-way ANOVA followed by Tukey’s multiple comparisons test (E)Septic shock/BSI with heat-killed cecal slurry rescued with *E. coli* BW25113 results in increased plasma putrescine metabolite levels; n = 5 HK CS, n = 9 HK CS + *E. coli* BW25113; Two-tailed p-values were determined by unpaired t test (F) *Klebsiella pneumoniae* pneumonia results in increased bronchial alveolar lavage (BAL) fluid levels of *N*- acetylputrescine; n = 4 PBS, n = 6 *K. pneumoniae* (KP9); Two-tailed p-values were determined by Mann Whitney test to include the outlier. (G) *Klebsiella pneumoniae* pneumonia results in increased plasma levels of *N*-acetylputrescine; n = 4 PBS, n = 6 *K. pneumoniae* (KP9); Two-tailed p-values were determined by Mann Whitney test to include the outlier. (H) Bacterial and mouse production of *N*-acetylputrescine + 4-acetamidobutanoate from putrescine; n = 3 for bacteria, representative data from 1-3 independent repeats; n = 6 mice; Blue = *E. coli*, Green = *K. pneumoniae*, Red = *P. aeruginosa* For all panels, data presented are means ± SEM; *p < 0.05; **p < 0.01; ***p < 0.001; ****p<0.0001

To further investigate whether putrescine and/or lysophosphatidyl choline metabolites were bacterially-derived, we compared plasma metabolomics profiles of sterile injury and live infection in mouse models of sepsis/BSI, hypothesizing that levels of microbial products would change only in the presence of live bacteria. Using the cecal-slurry (CS) model^30^, we compared an intraperitoneal (IP) injection of PBS, heat-killed CS (HK CS), and live CS and observed significant elevations in both *N*-acetylputrescine and 4-acetamidobutanoate only in those mice receiving live CS (**Figure 1D** and **Figure S2A-B**). By contrast, lysophosphatidyl cholines decreased in both HK and live CS (**Figure S2C**), consistent with a host-derived metabolic response to an inflammatory stimulus. Elevations in *N*-acetylputrescine and 4-acetamidobutanoate were also seen in the cecal ligation and puncture (CLP) model^31^ in the context of live infection, but not when utilizing *E. coli* lipopolysaccharide (LPS) as a sterile, inflammatory insult (**Figure S2D**). Thus in both models, plasma *N*-acetylputrescine and 4- acetamidobutanoate increases required a live infection, consistent with bacterial production of these metabolites. As both models involve polymicrobial infections, we also investigated whether these putrescine metabolites were altered in the context of single organism infections. Production of putrescine metabolites in HK CS was rescued by the addition of live *E. coli* (**Figure 1E**). A separate *Klebsiella pneumoniae* pneumonia^32,33^ model produced increased levels of putrescine metabolites (**Figure 1F-G** and **Figure S3**).

To confirm the microbial origin of putrescine-derived metabolites identified in our human and mouse studies, we cultured a panel of multidrug resistant (MDR) clinical isolates of *E. coli, K. pneumoniae,* and *P. aeruginosa* in minimal medium supplemented with putrescine and measured production of *N*-acetylputrescine and 4- acetamidobutanoate in culture supernatants. All strains tested produced *N*-acetylputrescine, while *P. aeruginosa* also produced the downstream metabolite 4-acetamidobutanoate (**Figure 1H** and **Figure S4A-B**)^26^. To assess production of these metabolites by mammalian hosts, we performed IP injections of putrescine in mice and measured plasma levels of *N*-acetylputrescine and 4-acetamidobutanoate (**Figure 1H** and **Figure S4A-E**). This demonstrated minimal generation of plasma putrescine metabolites in mammals, which is consistent with the previously reported restriction of mammalian *N*-acetylputrescine production to the brain, likely as a precursor for GABA synthesis^27,34^. Interestingly, only mice and *P. aeruginosa* produced 4- acetamidobutanoate from *N*-acetylputrescine whereas the remaining bacteria did not (**Figure S4F-H)**^34^, suggesting cooperative metabolism may explain some of the observed plasma metabolite alterations in both patients and mice with certain infections. Overall, these data highlight putrescine acetylation as a prominent microbial metabolic activity detectable in gram-negative BSI.

### SpeG homologs produce *N*-acetylputrescine in gram-negative pathogens

As a first step in investigating the importance of putrescine acetylation during BSI, we sought to identify the bacterial enzyme(s) responsible for this activity. Putrescine acetylation by *E. coli* has been reported previously^26^, but the responsible enzyme has not been identified. Polyamine/diamine *N*-acetyltransferases (EC: 2.3.1.57) quickly emerged as likely candidates. These enzymes are structurally diverse, and their activity has been biochemically validated across kingdoms^35^. *E. coli* and *K. pneumoniae* strains universally encode one member of this family, SpeG (sharing >89% amino acid ID), which is annotated as the putative putrescine *N*- acetyltransferase. However, its acetylation activity *in vitro* and in culture has only been demonstrated experimentally toward the longer chain polyamines spermidine and spermine, with acetylation of putrescine explicitly not observed *in vitro* or missed in culture^35-37^, possibly because the physiologically relevant substrate concentrations were not studied. This has led to the perception that spermidine is the substrate for SpeG despite it being found at 5-to-10-fold lower intracellular concentrations than putrescine in these microbes (1–5 mM compared with 20–30 mM). Spermine is not present in these bacteria^38-41^.

We investigated whether SpeG was responsible for putrescine acetylation in *E. coli*. By examining an *E. coli* BW25113 *speG* mutant previously generated as part of the KEIO collection^42^, we confirmed that deletion of *speG* resulted in a loss of *N*-acetylputrescine production in culture (**Figure 2A**). Complementation with a plasmid encoding *speG* rescued *N*-acetylputrescine production (**Figure 2A**), and incubation of purified enzyme with putrescine and acetyl-CoA established that SpeG catalyzes this reaction (**Figure 2B** and **Figures S5A**). We next sought to determine the kinetic parameters of SpeG toward putrescine to gain insight into why this activity had been missed previously. SpeG displayed activity toward putrescine at the high end of physiologic intracellular putrescine levels in *E. coli* (*K*half = 51.0 ± 5.4 mM) (**Figure 2C**)^38,40,43^. Although physiologic, the putrescine concentrations we used are higher than those previously examined, indicating how this activity could have escaped detection. As a control, we confirmed SpeG could acetylate spermidine and observed kinetic parameters similar to those published previously (**Figure S5B**)^35^. The *K*half (590.1 ± 30.8 µM) towards spermidine was well below the intracellular concentrations (1–5 mM)^38,40^, indicating that SpeG already operates near or at maximum velocity on spermidine under physiologic conditions. Consistent with SpeG’s dodecameric quarternary structure and allosteric binding sites, we observed cooperative kinetics with both putrescine and spermidine (**Figure 2C** and **Figure S5B**)^35,44^. This combination of kinetic parameters that reflect metabolite concentrations and cooperative kinetics suggests a previously overlooked role for SpeG as particularly responsive modulator of intracellular putrescine levels, enabling a rapid increase in activity when high concentrations are reached.

**Figure 2.**
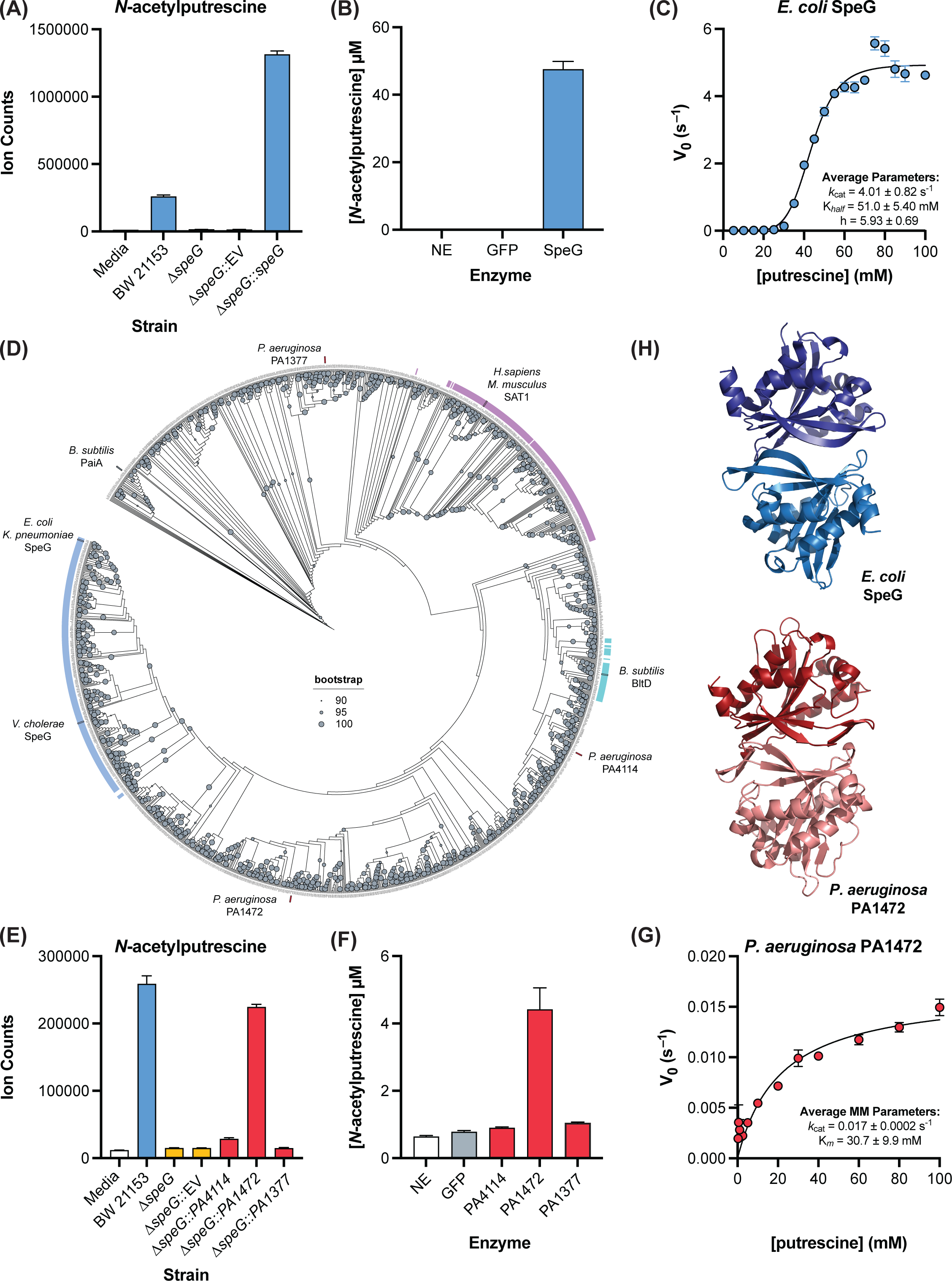
SpeG homologs are responsible for production of *N*-acetylputrescine in gram-negative pathogens. (A) Complementation in *E. coli* BW25113 demonstrates SpeG can produce *N*-acetylputrescine; n = 3 per condition, representative data from 3 independent experiments (B) 1 hour production of *N*-acetylputrescine from putrescine by recombinant purified enzymes; n = 3 per condition, representative data from 3 independent experiments; NE = no enzyme, GFP = green fluorescent protein (C) Kinetics of putrescine acetylation by SpeG is consistent with previously demonstrated cooperative mechanism on spermidine, n = 2-3 per substrate concentration, representative data from 4 independent experiments; summary parameters includes all experiments (D) Maximum-likelihood phylogenetic tree of a representative member of each group of protein sequences sharing >80% amino acid ID (RepNode80); Blue = *E. coli* SpeG clade, Purple = mammalian SAT1 clade, Turquoise = *B. subtilis* BltD clade (E) Complementation in *E. coli* BW25113 demonstrates PA1472 can produce *N*-acetylputrescine; n = 3 per condition, representative data from 2-3 independent experiments (F) 1 hour production of *N*-acetylputrescine from putrescine by recombinant purified enzymes; n = 3 per condition, representative data from 2-3 independent experiments; NE = no enzyme, GFP = green fluorescent protein (G) Kinetics of PA1472 on putrescine; n = 2 per concentration, representative data from 3 independent experiments; summary parameters includes all experiments (H) SpeG (PDB: 3WR7) and PA1472 (AlphaFold2) dimers For all panels, data presented are means ± SEM

Although patients with *P. aeruginosa* infection were included in our original cohort demonstrating an association between *N*-acetylputrescine and BSI, and we observed *P. aeruginosa* production of *N*-acetylputrescine in culture, *Pseudomonas* spp. lack a biochemically validated polyamine *N*-acetyltransferase. To begin identifying the *P. aeruginosa* enzyme(s) responsible for this metabolism, we searched the list of all putative polyamine/diamine *N*-acetyltransferases in the UniProt database, which are designated by EC 2.3.1.57^45,46^. This list included a single enzyme from *P. aeruginosa* PAO1, PA4114, which is annotated as a spermidine acetyltransferase sharing homology to the experimentally validated *Bacillus subtilis* BltD enzyme^47,48^. Additionally, we conducted a BLAST search using the *E. coli* SpeG amino acid sequence as a query and identified two additional candidate uncharacterized *P. aeruginosa* PAO1 enzymes, PA1377 and PA1472, which display low sequence identities to SpeG (26% amino acid ID each). We condensed the list of sequences from EC 2.3.1.57 into representative sequences sharing >80% amino acid sequence ID (i.e. RepNode80) and then performed a MUSCLE alignment^49^ with these sequences and the candidate *P. aeruginosa* sequences, followed by a maximum likelihood phylogenetic analysis. Strikingly, each of the three candidate *P. aeruginosa* enzymes clustered in clades containing a structurally distinct, validated polyamine acetyltransferase (**Figure 2D**)^50,51^.

To experimentally determine whether any of these candidate *P. aeruginosa* enzymes could acetylate putrescine, we complemented the *E. coli* BW25113 *speG* mutant^42^ with plasmids encoding *PA4114, PA1472*, and *PA1377.* This approach demonstrated that PA1472, but not PA4114 and PA1377, could rescue putrescine acetylation (**Figure 2E** and **Figure S5D**). Incubation of purified enzymes with putrescine and acetyl-CoA further established PA1472 as the likely *P. aeruginosa* SpeG homolog (**Figure 2F** and **Figure S5A**). We next determined the kinetic properties of PA1472 toward putrescine and confirmed that like SpeG, it is most active at the high end of intracellular putrescine concentrations (*Km* = 30.7 ± 9.9 mM) (**Figure 2G**)^41^. Consistent with PA1472 possessing putrescine *N*-acetyltransferase activity, its dimer AlphaFold2 predicted structure resembled the crystal structure of SpeG (3WR7, RMSD 1.891), despite the low % amino acid ID shared between the primary sequences (**Figure 2H**)^44^. Together, these data implicate PA1472, and not the annotated PA4114, as the likely putrescine *N*- acetyltransferase in *P. aeruginosa*. This finding highlights both the diversity of the enzymes capable of carrying out this chemistry and emphasizes the importance of experimental validation of computation hits when assigning enzyme functions.

Our initial sequential metabolomics studies on human samples, mouse models, and bacterial cultures highlighted putrescine acetylation as a microbially-mediated transformation detectable in BSI. To further confirm our assignment of *N*-acetylputrescine as bacterial in origin, we determined the kinetic parameters of the human homolog of SpeG, spermidine/spermine acetyltransferase 1 (SAT1), against putrescine. This revealed that SAT1 had a *K*m (8.70 ± 2.43 mM) well above reported intracellular putrescine concentrations in eukaryotic cells (sub-mM range). This is consistent with our mouse experiment (**Figure 1H**) indicating SAT1 contributes minimally to putrescine acetylation under physiologic conditions (**Figure S5A** and **Figure S5C**)^38^. As a control, we confirmed SAT1’s robust activity toward spermidine (*V/K* = 2.1 x 10^6^ M^-1^ min^-1^) (**Figure S5A** and **Figure S5D**)^52^. There was also little predicted structural similarity of either SpeG or PA1472 to SAT1 (2B5G, RMSD 11.454 and 12.187 respectively) (**Figure S5E**)^53^. Overall, these data confirmed the ability of SpeG to acetylate putrescine in *E. coli*, identified a likely *P. aeruginosa* putrescine acetyltransferase, and further suggested that bacterial enzymes are responsible for the increases in plasma *N*-acetylputrescine seen in BSI.

### Loss of SpeG activity impairs *E. coli* fitness in culture and *in vivo*

Across kingdoms, polyamines play diverse roles in the regulation of gene expression and protein translation and are implicated in the maintenance of cell membrane homeostasis and protection from oxidative stress^54^. As high levels of these polycationic molecules can be toxic^37^, intracellular concentrations must be tightly regulated, with charge neutralization through acetylation by SpeG thought to play a key role in bacteria^55^. *SpeG* deletion has been reported to hinder intracellular proliferation of *Salmonella*^56^ and to impair the resistance of the methicillin-resistant *Staphylococcus aureus* (MRSA) USA300 strain to extracellular host polyamines^57,58^. The importance of *speG* for pathogenesis may not be universal, however, as *Shigella* spp. have silenced *speG* expression to promote spermidine accumulation ^59^. Nevertheless, the increased levels of acetylated putrescine metabolites we observed in patient plasma suggests this activity may be important for the pathogens studied here, especially during the metabolically stressful process of BSI^15^.

To examine whether *speG* deletion is similarly important in *E. coli*, we compared growth of wildtype, *ΔspeG* and complemented BW25113 strains, which revealed no difference in proliferation (**Figure S6A**). To see if this response was strain-specific, we attempted to delete *speG* from two clinical *E. coli* isolates (M 1/5 and E23), but were unsuccessful, raising the possibility that *speG* might be essential in some strains and/or that BW25113 *ΔspeG* possessed an unappreciated compensatory change. To avoid both of these potential issues, we turned to an inducible CRISPRi system^60^, which resulted in ∼50% repression of *speG* transcription with a guide RNA targeting *speG* compared with a control guide targeting *RFP* in BW25113 (**Figure S6B**). This transcriptional repression resulted in a specific, dramatic suppression of *E. coli* proliferation in BW25113 (**Figure S6C**), calling into question the reliability of this deletion strain to elucidate secondary phenotypes related to *speG* loss. To address this concern, we then applied this system to the clinical *E. coli* isolates described above and observed closer to 70% repression of *speG* transcription (**Figure 3A** and **Figure S6D**). Consistent with a role in putrescine acetylation, *speG* repression resulted in significant decreases in extracellular *N*-acetylputrescine production (**Figure 3B** and **Figure S6E**), as well as significantly increased intracellular putrescine accumulation with a concomitant decrease in *N*-acetylputrescine (**Figure 3C** and **Figure S6F**). Reducing *speG* expression in these clinical isolates also recapitulated the growth defect observed with CRISPRi in *E. coli* BW25113 (**Figure 3D** and **Figure S6G-H**). Supplementation of the culture media with *N*-acetylputrescine did not rescue the proliferation defect, suggesting that SpeG’s regulation of polyamines, rather than the acetylated products themselves, conferred the proliferative advantage (**Figure S6I**). Finally, we sought to determine the consequences of reduction of *speG* expression *in vivo*. Using a modification of the rescue experiment from **Figure 1E**, we found that repression of *speG* transcription using inducible CRISPRi in *E. coli* E23 resulted in a significant delay in mortality compared with controls (**Figure 3E**). Combined with our earlier data, these findings support the hypothesis that increases in plasma *N*-acetylputrescine during bacterial infection arise from bacterial metabolism and that this highlights a metabolic process, polyamine acetylation, that plays an important role in pathogen physiology.

**Figure 3.**
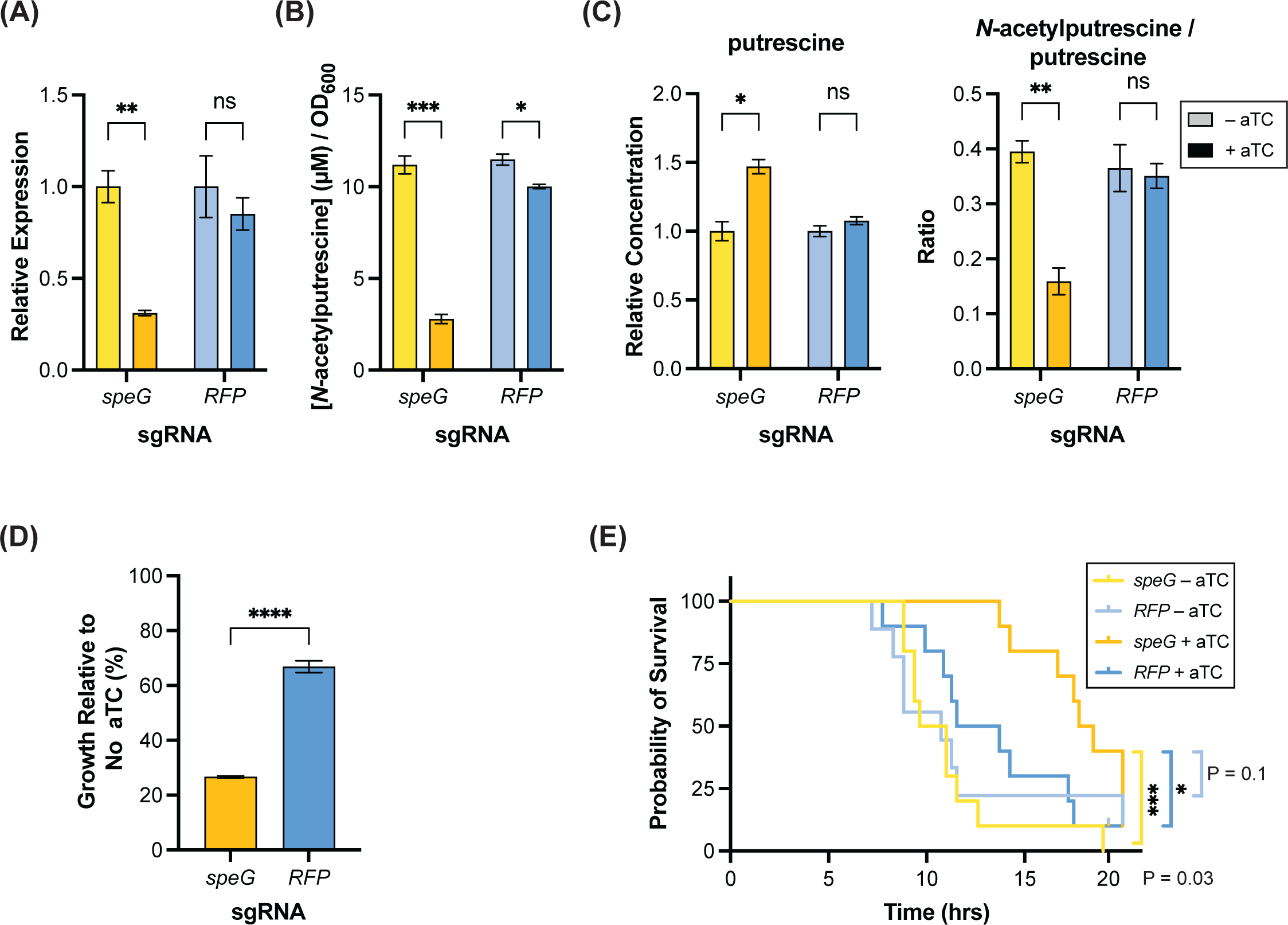
Suppression of *speG* expression impacts proliferation. (A) Suppression of *speG* expression with inducible CRISPRi in *E. coli* patient bloodstream isolate E23; n = 3 per condition, representative data from 2 independent experiments (B) Decreased levels of extracellular *N*-acetylputrescine with inducible CRISPRi of *speG* in E23, concentrations normalized to OD600 = 1.0; n = 3 per condition, representative data from 2 independent experiments (C) Increased relative intracellular putrescine levels (concentrations normalized to OD600 = 1.0 and then normalized to concentration of control condition) and decreased ratio of *N*-acetylputrescine/putrescine with inducible CRISPRi of *speG* in E23; n = 3 per condition, representative data from 2 independent experiments (D) Inducible CRISPRi of *speG* suppresses E23 growth in culture; n = 3 per condition, representative data from 3 independent experiments (E) Inducible CRISPRi of *speG* delays mortality in a cecal slurry model of sepsis with E23; n = 10 mice per group for all groups except n = 9 for *RFP* – aTC; p-value determined by Log-rank (Mantel-Cox) test For panels A-D, data presented are means ± SEM; Two-tailed p-values were determined by unpaired t test; *p < 0.05; **p < 0.01; ***p < 0.001; ****p<0.0001 aTC = anhydrotetracycline

### Diminazene inhibits SpeG and reduces bacterial proliferation

Given these observations, we sought a chemical method of inhibiting bacterial polyamine acetylation. While small molecule inhibitors of SpeG have not been reported, inhibitors of the human polyamine *N*- acetyltransferase SAT1 have been developed and studied as potential anti-cancer therapies^61,62^. It has also been previously shown that diminazene, a structural mimic of spermidine used a veterinary antitrypanosomal agent (**Figure 4A**), is an inhibitor of SAT1 with a *K*i of 2.4 µM^44,55,63^. Indirect evidence has suggested diminazene may inhibit the SpeG homolog from MRSA USA300, although it has not been tested against this enzyme directly^64^. To test if diminazene inhibits putrescine acetyltransferase activity of *E. coli* SpeG we determined IC50s for this drug *in vitro* using both putrescine and spermidine as substrates (**Figure 4B** and **Figure S7A**). Diminazene’s IC50 against SpeG with putrescine as a substrate (26.2 ± 3.7 µM) was similar to our measurement of its IC50 for SAT1 with spermidine (21.0 ± 0.6 µM) (**Figure S7B**), consistent with this drug being a broad-spectrum polyamine acetyltransferase inhibitor. In agreement with our CRISPRi experiments, diminazene treatment with doses similar to those used against MRSA USA300^64^ significantly decreased extracellular levels of *N*-acetylputrescine (**Figure 4C** and **Figure S7C**) and significantly increased intracellular putrescine accumulation while decreasing *N*-acetylputrescine levels (**Figure 4D** and **Figure S7D**) in clinical isolates. Diminazene treatment also inhibited growth in a dose-dependent manner (**Figure 4E** and **Figure S7E**) and increased sensitivity to drug treatment was observed during growth in minimal medium (**Figure S7E-G**).

**Figure 4.**
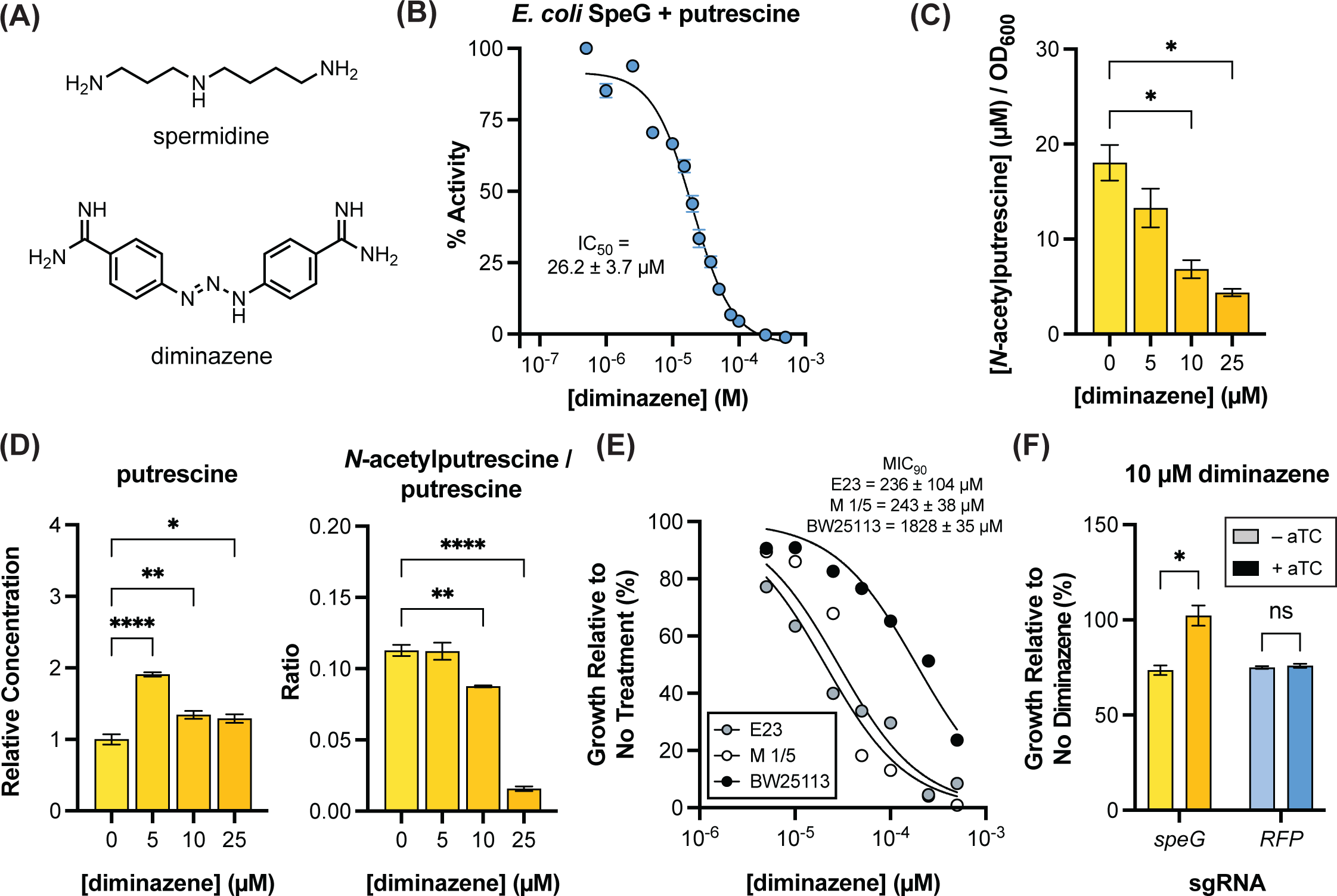
Diminazene inhibits SpeG and phenocopies inducible CRISPRi. (A) Structures of spermidine and diminazene (B) Determination of *in vitro* IC50 of diminazene against speG with putrescine substrate; n = 2 per inhibitor concentration, representative data from 3 independent experiments; summary data includes all experiments (C) Diminazene treatment reduces extracellular levels of *N*-acetylputrescine in E23, concentrations normalized to OD600 = 1.0; n = 3 per condition, representative data from 3 independent experiments; p-values were determined by One-way ANOVA followed by Dunnett’s multiple comparisons test with all comparisons made against no drug control (D) Diminazene treatment increases relative intracellular putrescine levels (concentrations normalized to OD600 = 1.0 and then normalized to concentration of control, 0 µM) condition) and decreases the ratio of *N*- acetylputrescine in E23; n = 3 per condition, p-values were determined by One-way ANOVA followed by Dunnett’s multiple comparisons test with all comparisons made against no drug control (E) MIC90 of diminazene treated E23, M 1/5, and BW25113 in LB; n = 3 per condition, representative data from 3 independent experiments; summary data includes all experiments (F) Inducible CRISPRi of *speG* blocks growth inhibition by diminazene in E23; n = 3 per condition, representative data from 2 independent experiments; Two-tailed p-values were determined by unpaired t test For all panels, data presented are means ± SEM; *p < 0.05; **p < 0.01; ***p < 0.001; ****p<0.0001 aTC = anhydrotetracycline

Given its structure and the pleiotropic roles of polyamines in cells, we wanted to test whether the growth suppression phenotype of diminazene in *E. coli* was SpeG-dependent or due to off-target effects. Prior work has suggested that diminazene may disrupt the proton motive force in *E. coli,* in addition to its known ability to bind nucleic acids^65^. There is also some evidence it could be broadly effective in *S. aureus* strains, including those likely lacking SpeG, albeit at higher doses^66^. Nevertheless, using our inducible CRISPRi system, we found that reducing *speG* expression rendered *E. coli* resistant to the growth inhibitory effects of 10 µM diminazene (**Figure 4F** and **Figure S7H**) indicating that at least at this modest dose, diminazene’s effects on bacterial growth are mediated through SpeG activity. Finally, we sought to determine MIC90s of diminazene across the previously studied MDR clinical isolates. Consistent with previous reports, diminazene sensitivity can vary across strains, with *E. coli* often more sensitive than *Klebsiella* spp. (**Figure S7I-J**)^66^. Together, these data suggest that we could leverage diminazene as a tool to further investigate the importance of polyamine acetylation across Gram-negative pathogens.

### Inhibiting polyamine acetylation increases bacterial membrane permeability and synergizes with clinical antibiotics

Several of the cellular processes regulated by polyamines^54^, which may be vulnerable to perturbations of polyamine metabolism, are also targets of antibiotics, with examples including DNA replication/transcription (quinolones), translation (macrolides and tetracyclines), and the cell membrane (beta-lactams and vancomycin)^15^. To elucidate which of these processes may be involved in the growth limitation phenotype observed upon repression of SpeG activity, we conducted drug synergy testing in the *E. coli* clinical isolates^67,68^ with diminazene and the antibiotics ciprofloxacin, erythromycin, tetracycline, and vancomycin. Using a checkerboard assay format, we found that diminazene synergized with vancomycin, a drug used exclusively to treat gram-positive infections (**Figure 5A** and **Figure S8A**), reducing the vancomycin MIC90 from above the limit of our assay (>128 µg/mL) to 32-64 µg/mL depending on the dose of diminazene. These findings suggest that SpeG inhibition by diminazene may be increasing permeability of the outer membrane, allowing this large glycopeptide drug, which would not normally permeate the outer membrane, to reach its target in the periplasm^69^. Importantly, inducible CRISPRi of *speG* produced a similar result, resulting in a clear reduction of the vancomycin MIC90 (**Figure 5B** and **Figure S8B-C)**.

**Figure 5.**
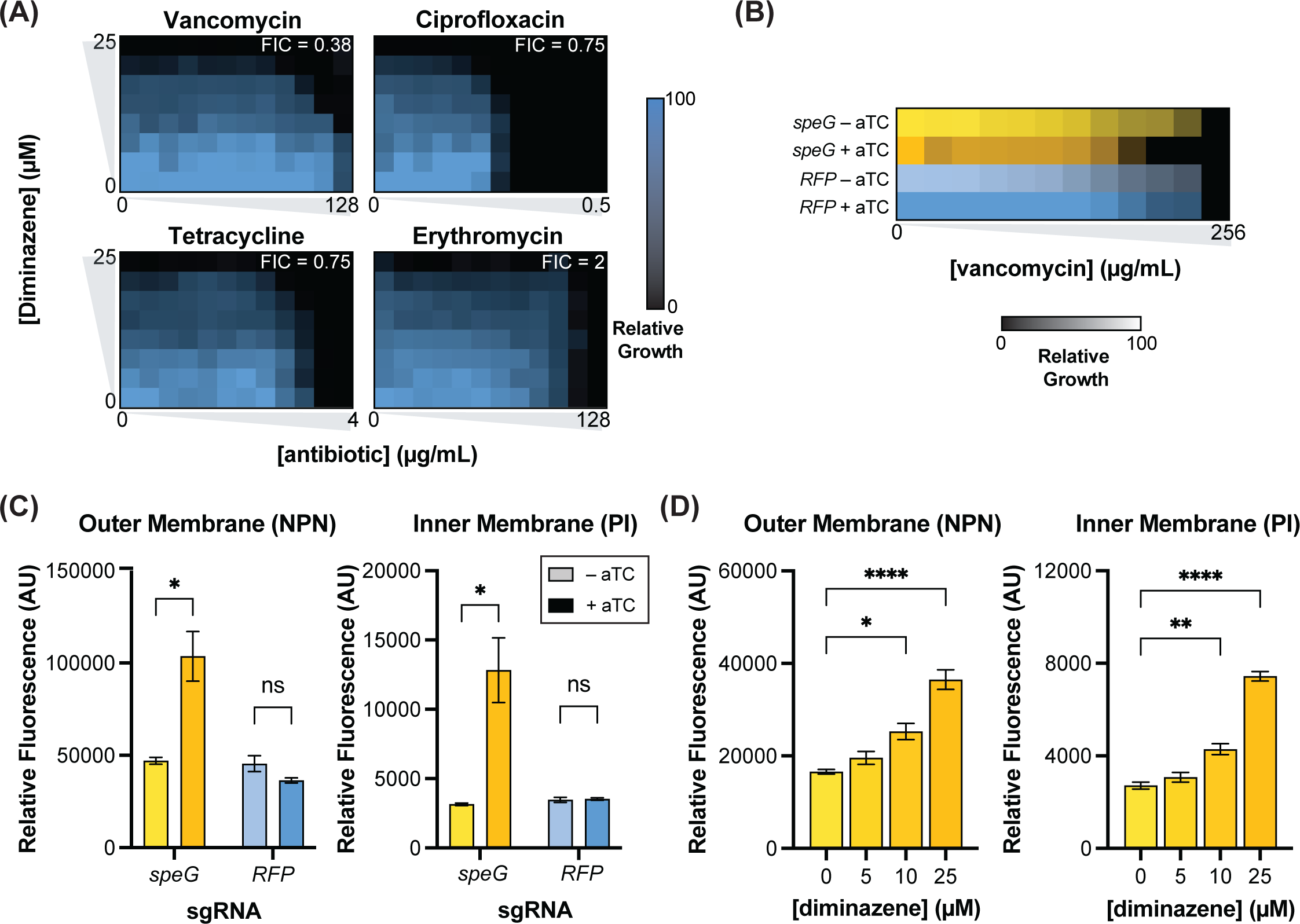
Reducing SpeG activity enhances membrane permeability. (A) Checkerboard assays demonstrate synergy between diminazene and vancomycin in *E. coli* E23; representative data from 3 independent experiments (B) Inducible CRISPRi of *speG* reduces MIC of vancomycin in *E. coli* E23; mean growth of n = 3 per condition, representative data from 3 independent experiments (C) Inducible CRISPRi of *speG* enhances membrane permeability in E23; n = 3 per condition, representative data from 2 independent experiments; Two-tailed p-values were determined by unpaired t test (D) Diminazene treatment for 6 hours enhances membrane permeability in *E. coli* E23; n = 3 per condition, representative data from 2 independent experiments; p-values were determined by One-way ANOVA followed by Dunnett’s multiple comparisons test with all comparisons made against no drug control For panels B-D, data presented are means ± SEM; *p < 0.05; **p < 0.01; ***p < 0.001; ****p<0.0001 aTC = anhydrotetracycline

We then directly assessed SpeG’s effect on bacterial inner and outer membrane permeability using a previously published fluorescence-based assay^70,71^. Inducible CRISPRi of *speG* increased permeability of both the outer and inner plasma membrane (**Figure 5C** and **Figure S8B**). Diminazene treatment for 6 hours phenocopied these observations in a dose-dependent manner (**Figure 5D** and **Figure S8E**), including at the dose at which we see SpeG-dependent growth inhibition (10 µM). Diminizene-mediated increases membrane permeability were greatly diminished with a shorter 3-hour treatment (**Figure S8F**), which is also consistent with SpeG inhibition driving this phenotype, rather than a direct effect on the outer membrane^69,72^. These data demonstrate that inhibiting the intracellular metabolic enzyme target SpeG can impact a membrane phenotype, creating a vulnerability that increases susceptibility to antibiotic treatment.

Two primary mechanisms of antibiotic resistance are limiting antibiotic entry into bacterial cells and efflux of antibiotics^73^. Given the effects of SpeG inhibition on membrane permeability, we hypothesized that diminazene treatment could perhaps re-sensitize MDR pathogens to existing antibiotics. Examining our panel of MDR patient isolates (**Figure 1H**), we specifically tested for synergy between diminazene and antibiotics to which these isolates were resistant. We found synergy between diminazene and tetracycline and/or ciprofloxacin in almost all these strains, in some cases re-achieving MIC90s at or below clinically relevant breakpoints (**Figure 6A** and **Figure S9A**). These findings were consistent with a recent report focused on repurposing old antimicrobials similar to pentamidine^69,72^, which found synergy between diminazene and streptomycin or chloramphenicol in some strains of drug resistant *E. coli*, *K. pnuemoniae*, and *S. aureus*, although the mechanism underlying this observation was not elucidated^66^. In our MDR strains, synergistic interactions were directly associated with increased intracellular accumulation of antibiotics (**Figure 6B** and **Figure S9B**) suggesting that diminazene treatment was improving cellular access of these drugs^74,75^.

**Figure 6.**
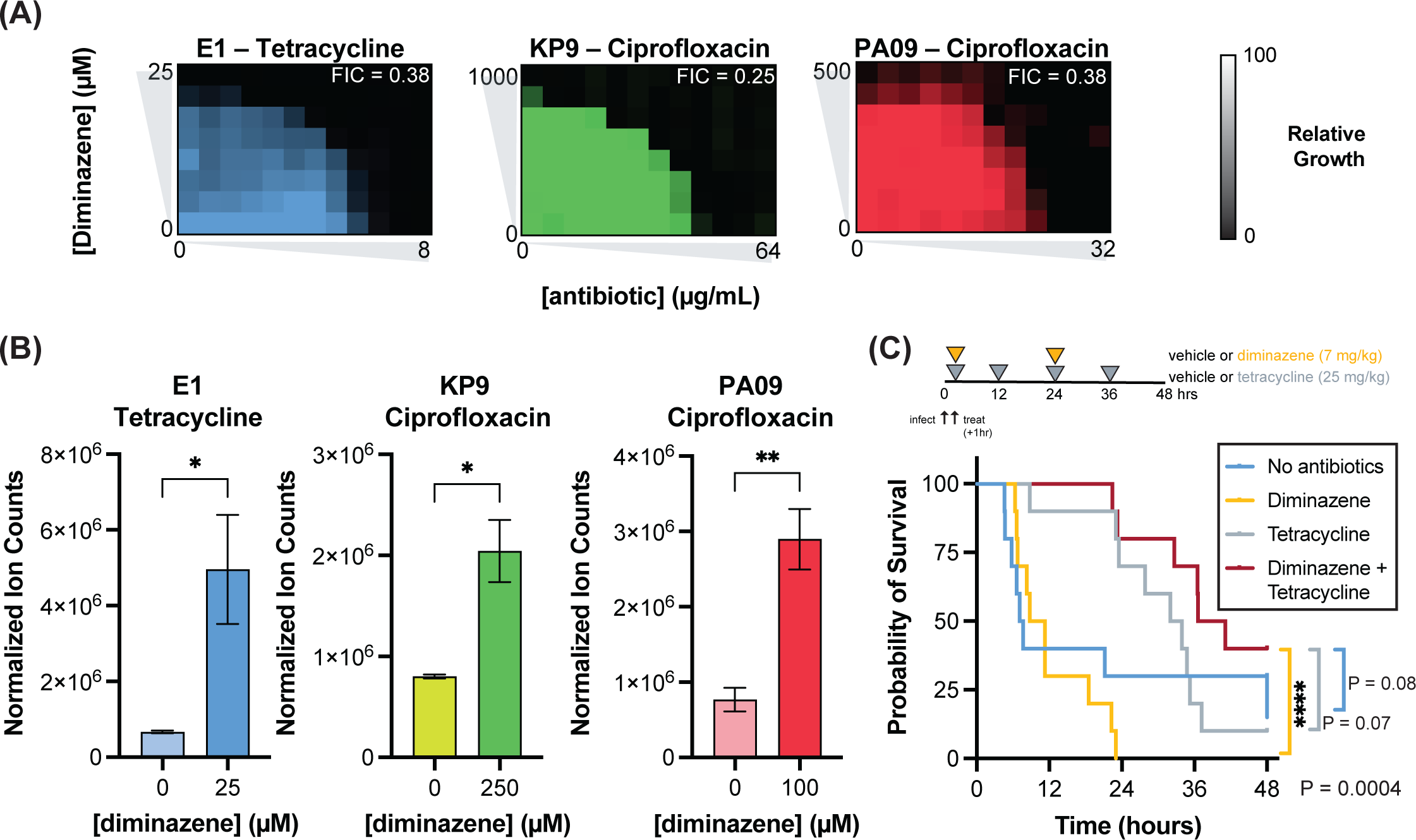
Blocking SpeG synergizes with existing clinical antibiotics in resistant bacteria in culture and*in vivo*. *(A)* Checkerboard assays demonstrate synergy between diminazene and antibiotics to which the assayed MDR strains are clinically resistant; representative data from 3-4 independent experiments per strain (B) Diminazene treatment enhances uptake of antibiotics to which MDR strains are resistant; n = 3 per condition, representative data from 1-3 independent experiments; Two-tailed p-values were determined by unpaired t test (C) Combination of diminazene and tetracycline treatment reduces mortality in a cecal slurry model of sepsis with the tetracycline resistant clinical isolate *E. coli* E1; n = 10 mice per group; p-value determined by Log-rank (Mantel-Cox) test For panels B, data presented are means ± SEM; *p < 0.05; **p < 0.01; ***p < 0.001; ****p<0.0001 aTC = anhydrotetracycline

Finally, we sought to test whether this synergy is observed *in vivo*. We focused on the tetracycline-resistant *E. coli* E1 strain, one of the strains for which diminazene treatment shifts the MIC90 back into the clinically sensitive range (**Figure 6A**). We performed a rescue of HK CS using the *E. coli* E1 line at a previously determined LD80-90 dose of bacteria. One hour after infection, we initiated treatment with clinically relevant doses of either vehicle, diminazene alone (7 mg/kg daily), tetracycline alone (25 mg/kg every 12 hours), or a combination of diminazene and tetracycline^76-80^. In vehicle and monotherapy groups, the expected mortality was observed by 48 hours, while combination therapy increased median and overall survival (**Figure 6C**). Tetracycline treatment alone improved median survival, but did not change the overall proportion of survivors (**Figure 6C**). This may indicate that peak concentrations of tetracycline after dosing could briefly overcome the resistance mechanism(s) utilized by E1, slowing but ultimately failing to clear the infection. Together, these data suggest that targeting SpeG activity therapeutically could provide opportunities to revisit our existing antibiotic arsenal and create new avenues for intervention in challenging to treat infections.

## DISCUSSION

Due to the rising rates of infections and deaths attributed to AMR pathogens, new and creative strategies are needed to replenish our dwindling pipeline of antibiotics^81^. Metabolism represents the biochemical manifestation of genetic, epigenetic, transcriptomic, and proteomic inputs which most accurately reflects an organism’s phenotype^82^. To this end, deciphering the metabolic activities of pathogens *in vivo* during infection is likely to highlight microbial pathways which support infection and could implicate new targets for intervention^83^. In this study, guided by our observation that *N*-acetylputrescine is elevated during BSI, we identify putrescine acetylation as a prominent microbial metabolic activity during infection and demonstrate that blocking this activity impairs pathogen fitness. This susceptibility occurred in part through increasing membrane permeability, which created an opportunity for synergy with existing clinical antibiotics. This strategy directly circumvents multiple modes of antibiotic resistance utilized by pathogens to limit intracellular access of drugs^73,81^. Thus, our workflow enables the identification of metabolic activities important for infection in patient samples. Characterizing the mechanism responsible for the observed changes and exploring its impact on pathogenesis ultimately allows us to leverage this understanding to identify new potential avenues for treatment.

The elevation of pathogen-derived *N*-acetylputrescine in human and mouse plasma during infection and its association with poor outcomes raises the possibility that this metabolite could be used clinically as marker of infections involving pathogens encoding SpeG homologs. Pathogen identification currently involves direct testing or culturing of biomaterial, which frequently requires several days^84^ and often misses a large proportion of BSIs^9^. The rapid identification of unique bacterial metabolites in patient biofluids could diagnose infection with a particular pathogen or group of pathogens possessing particular metabolic enzyme(s) independent of culture while simultaneously providing the guidance needed to specifically target the active metabolic pathways therapeutically. The presence or absence of bacterial metabolites could also be used to help guide antibiotic stewardship and limit unnecessary use of antibiotics, a known contributor to the development of AMR^4,85^. Supporting the potential of this approach, we found elevation of *N*-acetylputrescine in a parallel independent cohort of patients with septic shock compared with outpatient controls (R Rogers, *submitted*). Interestingly, this metabolite also distinguished patients between patients with septic shock and those with non-infectious, cardiogenic shock. Moving forward, larger validation cohorts are needed to determine formal characteristics and potential clinical utility of such a test.

Polyamines are ubiquitous across all life forms although the predominant intracellular polyamine molecules can vary widely across and within kingdoms^38,86^. They impinge on a multitude of cellular processes including the regulation of gene expression and protein translation, protection from oxidative stress, cell membrane function, cell growth, and pH tolerance^38,43,54,61,87-89^. At a chemical level, the amine groups of polyamines are protonated and positively charged at physiologic pH, while the alkyl backbones linking these amines enable hydrophobic interactions and impart flexibility^90,91^, such that the polycationic polyamines can bind to negatively charged macromolecules including DNA, RNA, membrane lipids, and proteins^38,40,43,90,91^. Given these promiscuous interactions, high concentrations of polyamines can be toxic^37^ and their levels are tightly regulated through a combination of synthesis, import/export, and metabolism^54^. Charge neutralization via acetylation of the terminal amines of the longer chain polyamines, spermidine and spermine, is thought to be a key metabolic mechanism of regulation, particularly under stressful conditions. In eukaryotic cells this reaction is performed by SAT1 while bacteria employ various acetyltransferases, including SpeG^55,92^. *SpeG* deletion has been reported to hinder intracellular proliferation of *Salmonella*^56^ and impair the resistance of the MRSA USA300 strain to host polyamines^57,58^, directly tying its activity to pathogen fitness. Nevertheless, the mechanism by which the loss of polyamine acetylation could impair fitness remained unclear.

Our identification of *N*-acetylputrescine as a bacterial metabolite elevated in BSI, and not *N*-acetylspermidine, was initially puzzling given previous reports of SpeG activity. To our knowledge, until this study there were no experimental data demonstrating SpeG’s activity toward putrescine. This is striking, as putrescine is the most abundant polyamine in the bacterial pathogens investigated here, while spermidine is present at about 5-to-10-fold lower concentrations^38-41^. Notably, under physiologic conditions, intracellular spermidine concentrations (1-5 mM) can be an order of magnitude greater than the corresponding *K*half for SpeG (∼600 µM), indicating that SpeG already operates on this substrate at or near its *V*max and perturbations in spermidine concentration are likely to have little impact on the rate of *N*-acetylspermidine production. Putrescine concentrations, in contrast, sit just below the measured *K*half (∼50 mM), indicating that SpeG likely plays a critical role in managing putrescine levels if they shift outside of tolerable ranges. To this end, SpeG’s cooperative kinetics^35,44^ enable a rapid and robust increase in activity under these conditions. Together, these observations indicate that, in addition to responding to spermidine and other exogenous polyamines, a major previously unrecognized role of SpeG and its homologs in the bacterial pathogens studied here is likely the management of intracellular putrescine levels.

Because polyamines interact with numerous cellular processes, we leveraged both genetic and chemical biology approaches to interrogate the impact of SpeG inhibition and prioritize pathways for further study. Our results highlighted membrane integrity as an SpeG-dependent property that may be exploited therapeutically, allowing us to significantly increase antibiotics’ access to cells and efficacy^93^. Our focus on membrane permeability, however, does not exclude the possibility that SpeG inhibition impacts other pathways within the cell. Further characterization of additional mechanism(s) by which SpeG inhibition impairs bacterial fitness are needed as disruptions of other pathways could provide important insight into SpeG’s role in enabling pathogenesis while also defining additional opportunities for intervention.

Due to the important role SpeG plays in the fitness of the gram-negative clinical isolates examined here, SpeG and its homologs represent an intriguing target across a broader range of AMR-associated pathogens. The remaining members of the top six AMR-associated pathogens^1^, *Acinetobacter baumannii*, *Streptococcus pneumoniae*, and *Staphyloccocus aureus*, all possess genes annotated as polyamine *N*-acetyltransferases (**Figure 2D**). The MRSA USA300 *speG* homolog is part of the Arginine Catabolic Mobile Element that has contributed to the clinical success of this strain and has been partially characterized with spermidine and spermine as substrates^57,58,64^. This homolog clusters with the *E. coli* and *K. pneumoniae* homologs in our phylogenetic analysis, confirming prior analyses showing a high degree of sequence similarity between these proteins^35^. Like *P*. *aeruginosa* PA1472, candidate homologs in *A. baumannii* and *S. pneumoniae* cluster separately, but map to the microbial side of the phylogenetic tree, distinct from eukaryotic homologs. Interestingly, a distinct polyamine acetyltransferase more similar to the mammalian SAT1 enzyme and not included in our phylogenetic analysis, was recently described in *A. baumannii,* with a narrow substrate scope predominantly limited to short chain 1,3-diaminopropane^94^. The *A. baumannii* candidate enzyme included in our phylogenetic analysis was not studied and its role in polyamine metabolism is unknown. Thus, while these initial leads are promising, further experimentation is needed to determine the functional roles of these candidate polyamine *N*-acetyltransferases and their potential contributions to pathogenesis.

While there were no demonstrated inhibitors of SpeG at the start of this study, we show that diminazene, a livestock antitrypanosomal drug and known inhibitor of SAT1, can also inhibit SpeG with similar potency^63,64^. Diminazene is not licensed for humans because of numerous toxicities in animals including convulsions, frequent urination and defecation, kidney and liver injury^95^. Despite being similarly susceptible to diminazene, the crystal structures of *E. coli* and *Vibrio cholerae* SpeG and *H. sapiens* SAT1 highlight several key structural distinctions, specifically a homo-dodecamer quaternary structure and the aforementioned allosteric polyamine binding site in the microbial enzyme compared with the simpler homodimer and single substrate binding site per subunit of the human enzyme^35,44,52^. These structural differences could provide an opportunity for the development of a microbial-specific polyamine acetyltransferase inhibitor, which could potentially target additional AMR-associated SpeG homologs. Interestingly, the recent solving of the MRSA USA300 SpeG homolog’s quaternary structure^96^ demonstrates marked overall similarity with the *E. coli* and *V. choerae* enzymes. Additionally, our AlphaFold2 prediction of PA1472 closely resembles *E. coli* SpeG, raising the possibility of a similar quaternary structure for this enzyme. Whether this is also true for the candidate *A. baumannii* and *S. pneumoniae* homologs and whether these structural predictions are experimentally supported will require further studies.

Overall, this study demonstrates the power of leveraging the natural, *in vivo* context of an infection to prioritize characterization of underappreciated aspects of microbial metabolism that contribute to disease. As our investigations of putrescine acetylation demonstrate, the resulting follow-up experiments can both inform our perspective on the key biological functions of specific microbial enzymes, identify new biomarkers that can guide therapy, and highlight potential vulnerabilities for therapeutic intervention. Thus, we anticipate that this study represents a starting point in developing a blueprint to investigate metabolism not only in gram-negative BSIs, but other infectious syndromes and clinically-relevant pathogens *in vivo*, with the ultimate goal of providing new tools to combat AMR.

## ACKNOWLEDGEMENTS

J.R.M., J.V., J.N. were supported by NHLBI T32 grant 5T32HL007633. M.D.I. was supported by a Canadian Institutes of Health Research Postdoctoral Fellowship (MFE-176575). N.R.G. was supported by an NSF Postdoctoral Research Fellowship in Biology under grant 1907240. R.M.B. was supported by R01 HL142093-01. E.P.B. was supported by the NSF Alan T. Waterman Award (CHE-20380529). E.P.B is a Howard Hughes Medical Institute Investigator.

## AUTHOR CONTRIBUTIONS

Conceptualization, J.R.M., R.M.B., and E.P.B.; Methodology, J.R.M., J.V., R.R.Z, M.D.I., A.B., N.R.G., J.N., C.H., M.A.P, C.B.C., S.D.Z., R.M.B., and E.P.B.; Investigation, J.R.M., J.V., R.R.Z., M.D.I., C.B., N.R.G., F.M.L,J.N, M.P.V., M.A.P., C.B.C., S.D.Z., R.M.B., and E.P.B.; Writing – Original Draft, J.R.M., R.M.B., and E.P.B;Funding Acquisition, R.M.B. and E.P.B.; Supervision, C.H., M.A.P. S.D.Z., R.M.B., and E.P.B.

## MATERIALS AND METHODS

### Human Metabolomics

#### Patient Samples

Patient samples were selected from the pre-existing Registry of Critical Illness (RoCI) cohort, housed at Brigham and Women’s Hospital (BWH) in Boston, MA, USA^97^. RoCI is approved by the Partners/Mass General Brigham Human Research Committee. Informed consent was obtained for blood collection. Curation of the database identified 21 patients with positive blood cultures growing either *Escherichia coli, Klebsiella pneumoniae,* or *Pseudomonas aeruginosa* with contemporaneous or near contemporaneous plasma samples banked and available for metabolomic analysis. Twenty-two controls admitted to the intensive care unit for reasons other than septic shock were identified for use as controls.

#### Metabolomics

Metabolomic measurements were made using 3 complementary liquid chromatography–tandem mass spectroscopy (LC-MS) methods. For each method, pooled plasma reference samples were included every 20 samples, and results were standardized using the ratio of the value of the sample to the value of the nearest pooled reference multiplied by the median of all reference values for the metabolite.

HILIC analyses of water-soluble metabolites in the positive ionization mode (HILIC-pos) were conducted using an LC-MS system composed of a Shimadzu Nexera X2 U-HPLC (Shimadzu Corp) coupled to a Q Exactive hybrid quadrupole orbitrap mass spectrometer (Thermo Fisher Scientific). Plasma samples (10 μL) were prepared via protein precipitation with the addition of 9 volumes of 74.9:24.9:0.2 v/v/v acetonitrile/methanol/formic acid containing stable isotope-labeled internal standards (valine-d8, Sigma-Aldrich; and phenylalanine-d8, Cambridge Isotope Laboratories). The samples were centrifuged (10 minutes, 9000*g*, 4 °C), and the supernatants were injected directly onto a 150×2 mm, 3-μm Atlantis HILIC column (Waters). The column was eluted isocratically at a flow rate of 250 μL/min with 5% mobile phase A (10 mmol/L ammonium formate and 0.1% formic acid in water) for 0.5 minute followed by a linear gradient to 40% mobile phase B (acetonitrile with 0.1% formic acid) over 10 minutes. Mass spectroscopic (MS) analyses were performed by using electrospray ionization in the positive ion mode using full scan analysis over 70 to 800 *m/z* at 70000 resolution and 3 Hz data acquisition rate. Other MS settings were as follows: sheath gas 40, sweep gas 2, spray voltage 3.5 kV, capillary temperature 350 °C, S-lens RF 40, heater temperature 300 °C, microscans 1, automatic gain control target 1e6, and maximum ion time 250 ms.

HILIC analyses of water-soluble metabolites in the negative ionization mode (HILIC-neg) were conducted by using an LC-MS system composed of an AQUITY UPLC system (Waters and a 5500 QTRAP mass spectrometer [SCIEX]). Plasma samples (30 μL) were prepared via protein precipitation with the addition of 4 volumes of 80% methanol containing inosine-^15^N4, thymine-d4, and glycocholate-d4 internal standards (Cambridge Isotope Laboratories). The samples were centrifuged (10 minutes, 9000*g*, 4 °C), and the supernatants were injected directly onto a 150×2.0 mm Luna NH2 column (Phenomenex). The column was eluted at a flow rate of 400 μL/min with initial conditions of 10% mobile phase A (20 mmol/L ammonium acetate and 20 mmol/L ammonium hydroxide in water) and 90% mobile phase B (10 mmol/L ammonium hydroxide in 75:25 v/v acetonitrile/methanol) followed by a 10-minute linear gradient to 100% mobile phase A. MS analyses were performed using electrospray ionization and selective multiple reaction monitoring scans in the negative ion mode. To create the method, declustering potentials and collision energies were optimized for each metabolite by infusion of reference standards. The ion spray voltage was –4.5 kV and the source temperature was 500 °C.

Positive ion mode analyses of polar and nonpolar plasma lipids (C8-pos) were conducted using an LC-MS system composed of a Shimadzu Nexera X2 U-HPLC (Shimadzu Corp) coupled to a Exactive Plus orbitrap mass spectrometer (Thermo Fisher Scientific). Plasma samples (10 μL) were extracted for lipid analyses using 190 μL of isopropanol containing 1,2-didodecanoyl-*sn*-glycero-3-phosphocholine (Avanti Polar Lipids). After centrifugation, supernatants were injected directly onto a 100×2.1 mm, 1.7-μm ACQUITY BEH C8 column (Waters). The column was eluted isocratically with 80% mobile phase A (95:5:0.1 v/v/v 10 mmol/L ammonium acetate/methanol/formic acid) for 1 minute followed by a linear gradient to 80% mobile phase B (99.9:0.1 v/v methanol/formic acid) over 2 minutes, a linear gradient to 100% mobile phase B over 7 minutes, then 3 minutes at 100% mobile phase B. MS analyses were performed using electrospray ionization in the positive ion mode using full scan analysis over 200 to 1000 *m/z* at 70000 resolution and 3 Hz data acquisition rate. Other MS settings were as follows: sheath gas 50, in source collision-induced dissociation 5 eV, sweep gas 5, spray voltage 3 kV, capillary temperature 300 °C, S-lens RF 60, heater temperature 300 °C, microscans 1, automatic gain control target 1e6, and maximum ion time 100 ms. Lipid identities were determined based on comparison with reference plasma extracts and were denoted by the total number of carbons in the lipid acyl chain(s) and total number of double bonds in the lipid acyl chain(s).

Raw data from Q Exactive/Exactive Plus instruments were processed using TraceFinder software (ThermoFisher Scientific) and Progenesis QI (Nonlinear Dynamics), whereas MultiQuant (SCIEX) was used to process 5500 QTRAP data. For each method, metabolite identities were confirmed using authentic reference standards or reference samples.

#### Statistical Analyses

##### Targeted metabolomics analyses

To normalize and standardize the LC–MS data, measurements from the internal standards were first corrected by removing the technical variation associated with injection order. This technical variation was characterized by regressing the internal standards readings on injection order using a local degree-two polynomial regression with an alpha parameter of 5, implemented using the R function loess. To correct for technical variation, internal standards were converted to ratios relative to the fitted readings from the regression. These ratios were then averaged across all internal standards used. The same procedure was used to correct for technical variation from each of the profiled metabolites. Finally, normalized metabolite readings were calculated by subtracting natural log-transformed average corrected standard from natural log-transformed corrected metabolite readings. Only metabolites that had less than 10% missing data in each of the two patient groups (septic shock and control patients) were considered.

To identify candidate metabolites, each metabolite was tested using a Wilcoxon to test to determine whether it was present at different levels between the two patient groups. Metabolites with false discovery rate-adjusted p-values less than 0.05 were deemed significantly differentially abundant. Each metabolite was also tested for association with APACHE II scores among all patients using Spearman’s correlation, and with mortality among all patients using a Wilcoxon test. A false discovery rate cutoff of 0.05 was used to identify statistically significant metabolites.

### Mouse Models

#### Models of Infection

All mouse experiments were conducted in accordance with protocols approved by the BWH Institutional Animal Care and Use Committee (IACUC). For most experiments, we used C57BL/6N SPF^98^ male mice 10-12 weeks of age purchased from Charles River Laboratories. The cecal slurry studies were also conducted in a cohort of C57BL/6N SPF female mice 10-12 weeks of age purchased from Charles River Laboratories. All mice were allowed to acclimate for more than one week prior to use.

##### Cecal Ligation and Puncture (CLP)

CLP was performed as previously described^99^. Briefly, after induction of anesthesia and analgesia with ketamine (85 mg/kg) and xylazine (34 mg/kg), mice were anesthetized, and a midline laparotomy was performed. The cecum was externalized, ligated, and punctured, after which a small amount of cecal contents were extruded from the puncture holes. The cecum was then replaced in the abdomen, and the abdominal incision was closed in layers. The mice were resuscitated with 1 mL of phosphate-buffered saline (PBS) and placed in a warmed cage for postoperative recovery. As our study was focused on identifying bacterial metabolites in plasma, the mice did not receive antibiotics to avoid reducing the sensitivity of our analyses. Mice were sacrificed at 24 hours. Original groups were n = 4 mice in PBS and CLP, n = 3 for LPS. One CLP mouse did not survive the full 24 hours and was excluded from analysis.

##### Cecal Slurry

Cecal slurry was performed as previously described^30^. Briefly, 10-14 week old C57B/6 mice from Charles River were sacrificed and whole cecum dissected. The entire cecal contents were collected with sterile forceps, spatula, and culture dishes. Collected contents were pooled and weighed before mixing 0.5 mL sterile water per 100 mg cecal contents. After resuspension, the cecal slurry was filtered through 100 µm sterile mesh filters (Falcon). The filtered cecal slurry was then mixed with an equal volume of sterile 30% glycerol in PBS. This final solution was aliquoted and stored at –80 °C until use.

Heat killed cecal slurry was produced by heating room temperature cecal slurry at 72 °C for 15 min. After cooling to 37 °C, an undiluted aliquot was plated on LB Lennox agar (VWR) to confirm the absence of culturable bacteria. For male mice, sepsis was induced with a 200 µL intraperitoneal injection of thawed live cecal slurry (live bacteria confirmed by plating). As controls, 200 µL of heat killed cecal slurry and 200 µL of 15% glycerol in sterile PBS were used. Mice were sacrificed at 24 hours. Original groups were n = 8, mice that did not survive the full 24 hours were excluded from analysis. For the smaller female mice, sepsis was induced with a 150 µL intraperitoneal injection of thawed live cecal slurry (live bacteria confirmed by plating). As controls, 150 µL of heat killed cecal slurry and 150 µL of 15% glycerol in sterile PBS were used. Female mice were sacrificed at 20 hours. Original groups were n = 6 each for PBS and heat-killed cecal slurry and n = 12 for live cecal slurry.

For the *E. coli* BW25113 HK CS rescue experiment, *E. coli* were grown overnight to stationary phase in LB Lenox Broth (VWR) before cells were washed 2x with PBS and OD600 determined by NanoDrop 2000c (Thermo Scientific). Cultures were spun down and resuspended to a calculated 1 x 10^8^ CFU/mL (http://www.agilent.com/store/biocalculators/calcODBacterial.jsp) in heat killed cecal slurry prior to administration. Actual CFU was determined to be ∼4 x 10^7^ CFU/mL by plating serial dilutions. 200 µL of this resuspension was injected into the peritoneal space of each mouse and mice were sacrificed at 24 hours. Groups were n = 5 heat-killed cecal slurry and n = 9 heat-killed cecal slurry + *E. coli* BW25113

For the CRISPRi experiment HK CS rescue experiment, mice received 3 days of 20 µm filter-sterilized drinking water ± 250 mg/L anhydrotetracycline (Cayman Chemicals) in red-tinted bottles (Ancare) to inhibit degradation by light. This was continued after administration of bacteria until the completion of the experiment. *E. coli* (E23) CRISPRi cell lines (details below) were grown overnight to stationary phase in LB Lenox Broth (VWR) before dilution 1:50 into fresh media in the morning ± 2 µM anhydrotetracycline. After 5 hours of culture, cells were washed 2x with PBS and OD600 determined by NanoDrop 2000c (Thermo Scientific). Cultures were spun down and resuspended to a calculated 5.0 x 10^7^ CFU/mL (http://www.agilent.com/store/biocalculators/calcODBacterial.jsp) in heat killed cecal slurry (± 2 µM anhydrotetracycline) prior to administration. Actual CFU was determined to be ∼2 x 10^7^ CFU/mL by plating serial dilutions. 200 µL of this resuspension was injected into the peritoneal space of each mouse and survival monitored over the proceeding 20 hours. Originally planned as n = 10 mice per group. One mouse planned for the *RFP* – aTC group was excluded prior to the start of the experiment for fight wounds.

For the antibiotic synergy experiment, the *E. coli E1* strain was grown overnight to stationary phase in LB Lenox Broth (VWR) before cells were washed 2x with PBS and OD600 determined by NanoDrop 2000c (Thermo Scientific). Cultures were spun down and resuspended to a calculated 5 x 10^9^ CFU/mL (http://www.agilent.com/store/biocalculators/calcODBacterial.jsp) in heat killed cecal slurry prior to administration. 200 µL of this resuspension was injected into the peritoneal space of each mouse. Actual CFU was determined to be ∼4 x 10^9^ CFU/mL by plating serial dilutions. Beginning one hour of after infection, mice received one of four treatments in equal volumes administered via IP injection: sterile 0.9% saline, diminazene in sterile 0.9% saline 7 mg/kg daily, tetracycline in sterile 0.9% saline 25 mg/kg every 12 hours, or both diminazene and tetracycline. Survival was monitored over the proceeding 48 hours.

##### Intranasal inoculation for pneumonia

Intranasal inoculation of *Klebsiella pneumonia* (KP9) was carried out as previously described^32,33^. Briefly, bacteria were grown to stationary phase overnight in LB Lennox Broth (VWR) before washing 2x with PBS and checking OD600 NanoDrop 2000c (Thermo Scientific). Cells were diluted to an OD600 = 0.8 for use. Actual CFUs were calculated to be ∼2.2 x 10^9^ CFU/mL by serial dilution. Mice were anesthetized as above with ketamine and xylazine and 40 µL of cells administered intranasally. Mice were kept vertical for ∼30-60 seconds to ensure delivery to the lungs prior to recovery from anesthesia. After 16 hours, mice were anesthetized again as above and their tracheas cannulated. Bronchial alveolar lavage with 1 mL of sterile PBS was performed (4 washes with same PBS solution), cells pelleted, and cell free supernatant was frozen at –80 °C for metabolomics analysis. Mice were subsequently sacrificed.

#### Putrescine and N-acetylputrescine injections

Mice received an intraperitoneal injection with either 100 mg/kg putrescine dihydrochloride (Sigma Aldrich), 50 mg/kg *N*-acetylputrescine hydrochloride (Sigma Aldrich) dissolved in PBS, or PBS as indicated. Area under the curve (AUC) was calculated for each individual mouse as µM*min (nmol*min/mL) and baseline (PBS injected average) AUC was subtracted. Estimated blood volume was calculated as 0.08 mL per g body weight^100,101^ and used determine total nmol of metabolite in blood over the course of the experiment. This was then normalized to estimated dry weight (26% of total weight)^102^ to determine nmol metabolite per g of dry weight.

#### Plasma samples

Plasma was collected for each experiment at the time points indicated. For repeat bleeding experiments, the tail vein was accessed and small amounts of blood harvested, in total reaching less than 10% of the total blood volume. At experimental completion, mice were anesthetized with ketamine and xylazine as above and terminally bled prior to sacrifice. In all cases, blood was placed in EDTA-pretreated tubes (Sarsedt) and centrifuged to separate plasma. Plasma was aliquoted and frozen at −80 °C for further analysis.

### Bacteria Experiments

#### Cell Lines

The *Escherichia coli* BW25113 parental strain and *ΔspeG* mutant (CGSC#: 9346, Name: JW1576-1) were obtained from the KEIO collection^42^ housed at Yale University (cgsc.biology.yale.edu/KeioList.php). *E. coli* M 1/5 strain is a human isolate that was a generous gift from Professor Ulrich Dobrindt (University of Münster). All remaining cell lines described are clinical patient isolates provided as a generous gift by Dr. Sophia Koo (Brigham and Women’s Hospital).

#### Putrescine derivative production

Indicated cell lines were growth overnight at 37 °C in M9 minimal media (1x M9 Salts, 0.1 mM CaCl2, 2 mM MgSO4, 0.4% glucose, 0.2% CAS amino acids, pH 7.3-7.4) to stationary phase in 5 mL culture. The following morning cells were pelleted and resuspended in 1 mL fresh M9 minimal media lacking CAS amino acids and supplemented with either 10 mM putrescine or *N*-acetylputrescine. Cells were then cultured for 5 hours at 37 °C. Half of the culture was applied to pre-weighed Whatman filter paper, dried overnight at 55 °C, and then filters re-weighed to determine dry cell mass. CFUs were calculated via serial dilution and plating on LB Lennox Agar. The remaining culture was pelleted and spent media removed and frozen at –80 °C for subsequent LC– MS analysis. Metabolite amounts were normalized to CFUs or dry mass as indicated.

#### Complementation

For all complementation studies, a truncated version of the pTrcHis 2A expression vector (ThermoFisher) was utilized, which allowed use of the native *E. coli* K12 *speG* promoter. All promoters were designed with the NEBuilder assembly tool (http://www.nebuilder.neb.com/-!/). To obtain the native *E. coli* sequence for use, *speG* and its promoter were amplified via PCR from genomic DNA harvested from *E. coli* K12 MG1655. After PCR amplification, this sequence was cloned into the truncated pTrcHis vector using Gibson assembly. After the correct sequence was confirmed with DNA sequencing, this vector was transformed into *E. coli* BW25113 *ΔspeG* using electroporation. This construct containing the *speG* promoter was then used to assemble of the remaining complementation constructs utilized. To generate these constructs, amino acid sequences were obtained from the UniProt database (http://uniprot.org) and uploaded to the ThermoFisher GeneArt synthesis portal, which generated *E. coli* codon-optimized versions of each gene. After PCR amplification, these sequences were cloned into the modified pTrcHis + *speG* promoter vector using Gibson assembly. As before, the correct sequence was confirmed with DNA sequencing prior to transformation into *E. coli* BW25113 *ΔspeG* using electroporation. All primers and codon optimized gene sequences used are listed in **Table S3–S4**.

To test for polyamine metabolite production, stationary phase cultures were diluted 1:1000 into LB Lennox and grown overnight at 37 °C. Parental BW25113 was grown without antibiotics, *ΔspeG* with Kanamycin (VWR, 50 µg/mL), and complemented strains with Kananmycin and Ampicillin (Sigma Aldrich, 100 µg/mL). The following morning OD600 was measured by plate reader (BioTek Synergy HTX), cells were pelleted, and spent media was frozen at –80 °C for subsequent analysis.

#### Growth curves

Stationary phase cultures were diluted 1:100 into either LB Lenox or M9 minimal media with 0.4% glucose and 0.2% CAS-amino acids as indicated. Cultures were performed in either 96 or 384-well plates, maintained at 37 °C and OD600 measured after shaking every 10 minutes for 16 hours by plate reader (BioTek Synergy HTX). For CRISPRi and MIC90 experiments, anhydrotetracycline (Cayman Chemical) or diminazene (Selleck Chemicals) respectively were added at the time of dilution. For growth relative to no treatment controls, the area under the curve (AUC) was calculated for each individual biological replicate of the experimental condition and normalized to the average of the no treatment control AUC.

#### CRISPRi

The inducible CRISPRi system consisting of pdCaS7-bacteria (AddGene #44249) and pgRNA-bacteria (AddGene #44251) was a gift from Stanley Qi ^60,103^. Guide sequence targeting *speG* was designed using Millepore Sigma’s CRISPR Design Tools. Cloning was performed as described in ^60^. Guide sequences for *E. coli* BW25113 and M 1/5 (5’->3’) ttaaaggtgattctacacca and for *E. coli* E23 (5’->3’) ttaaagctgattctacacca. Doses of the pdCaS7 expression inducer anhydrotetracyline (Cayman Chemical) used were 2 µM for BW25113, 0.5 µM for M 1/5, and 2 µM for E23. For CRISPRi + 10 µM diminazene experiment, 0.25 µM and 1.5 µM concentrations of anhydrotetracyline were used for M 1/5 and E23 respectively. All cultures were grown in media containing Ampicillin (Sigma Aldrich, 100 µg/mL) and chloramphenicol (Sigma Aldrich, 25 µg/mL). For gene expression, membrane permeability, and metabolite production experiments, cells were diluted 1:50 into fresh media supplemented with anhydrotetracycline. Growth curves were diluted 1:100 as discussed above. Transcriptional repression was assessed after 6 hours of induction, growth curves and membrane permeability after 16 hours of induction, and polyamine metabolite production with 6 hours of induction, followed by repeat dilution 1:50 and additional 16 hours of induction.

#### RT-qPCR

Cells pellets were resuspended in Direct-zol (Zymo) before RNA isolation using the Direct-zol RNA Miniprep Kit (Zymo). RNA quality and concentration was checked using Nano-drop 2000 (Thermo Scientific) with a target concentration of ∼10 ng/µL. RT-qPCR was then performed using Luna Universal One-step RT-qPCR kit (New England Biolabs) following the provided protocol on a BioRad CFX Opus 96 quantitative Real-Time PCR machine. Primers were previously published^104-106^ (**Table S5**) and efficiency of all the primers used was confirmed to be 95 ± 5% using a standard curve of template. Melt curves were performed to confirm the absence of primer dimers and the presence of a single amplicon. Samples were normalized to the geometric mean of a panel of endogenous control genes as previously described^107,108^.

#### Intracellular Putrescine Accumulation and Extracellular Production

For CRISPRi experiments, the stationary phase cultures of the indicated cell lines were diluted into 1:50 fresh LB Lennox media ± anhydrotetracycline (0.5 µM for M 1/5 and 2 µM for E23) and cultured for 5-6 hours shaking at 37 °C, before repeat dilution at 1:50 into fresh LB Lennox media ± anhydrotetracycline and overnight culture for 16 hours shaking at 37 °C. The following morning, OD600 was measured and cells pelleted. Spent media was removed and frozen at –80 °C for subsequent LC–MS analysis. Cell pellets were harvested as below.

For diminazene experiments, stationary phase cultures of the indicated cell lines were diluted 1:50 into fresh LB Lennox media and cultured for 5-6 hours shaking at 37 °C, before repeat dilution at 1:50 into fresh LB Lennox media with the indicated concentrations of diminazene and overnight culture for 16 hours shaking at 37 °C. The following morning, OD600 was measured and cells pelleted. Spent media was removed and frozen at –80 °C for subsequent LC–MS analysis. Cell pellets were harvested as below.

Cell pellets were washed twice with PBS and then resuspended in LC–MS grade water (400 µL per 1 mL culture). Resuspended cells underwent two freeze thaw cycles alternating between liquid nitrogen and heating to 65 °C. Cell debris was pelleted and 90% of aqueous supernatant removed. LC–MS grade methanol (200 µL per 1 mL culture) was added to the cell debris. Cell debris was re-pelleted and methanol supernatant combined with the previously removed aqueous supernatant. This 2:1 water:methanol mix was centrifuged one additional time and the resulting supernatant was ready for LC–MS analysis (described below).

#### Antibiotic synergy

Stationary phase cultures of indicated bacterial strains were diluted 1:5000 into M9 minimal media + 0.4% glucose + 0.2% CAS-amino acids containing indicated combinations of diminazene (Cayman Chemical) and vancomycin HCl (Research Products International), ciprofloxacin (Sigma Aldrich), tetracycline (Sigma), or erythromycin (Sigma). Drug concentrations were serially diluted 50% at each increment from the top indicated dose and checkerboard pattern was prepared by Tecan D300e Droplet Dispenser. Cells were cultured shaking overnight at 37 °C and OD600 measured the following morning by plate reader (BioTek Synergy Neo2). Background was subtracted from each well and then each well was normalized to the OD600 of the well without drug to calculate relative growth. MIC90 for each drug was determined to be the column or row in which there was no growth seen. In cases where growth was seen in every column or row, MIC90 was estimated to be 2x the highest concentration of drug. Fractional inhibitory concentration (FIC) was calculated FIC = FICA + FICB = (CA/MICA) + (CB/MICB), where MICA and MICB are the MIC90s of drugs A and B alone, respectively, and CA and CB are the concentrations of the drugs in combination corresponding to an MIC. We report here the FICmin from multiple repeat experiments. Synergistic interactions were defined as FIC <0.5, additive/indifferent interactions were defined as 0.5≤ FIC ≤4, and antagonistic interactions were defined as FIC >4^67,68^.

#### Determination of vancomycin MIC with CRISPRi

Stationary phase cultures of indicated cell lines were diluted 1:1000 into LB media LB Lennox media ± anhydrotetracycline (0.25 µM for M 1/5 and 1.5 µM for E23) along with vancomycin serially diluted at each increment from the top dose (256 µg/mL) as prepared by Tecan D300e Droplet Dispenser. Cells were cultured shaking overnight at 37 °C and OD600 measured the following morning. Background was subtracted from each well and then each well was normalized to the OD600 of the well without vancomycin.

#### Membrane Permeability

For CRISPRi experiments, the stationary phase cultures of the indicated cell lines were diluted 1:50 into fresh LB Lennox media ± anhydrotetracycline (0.5 µM for M 1/5 and 2 µM for E23) and cultured overnight for 16 hours shaking at 37 °C. For diminazene experiments, stationary phase cultures of the indicated cell lines were diluted 1:50 into fresh LB Lennox media and cultured for either 3 or 6 hours shaking at 37 °C. Assessment of inner and outer membrane permeability was adapted previously published protocols^70,71^. Briefly, cells were pelleted and washed 3x with 5 mM HEPES and 5 mM glucose buffer (pH = 7.2) before being resuspended in the same before and OD600 measured. Resuspended cells were split and incubated with either 1-*N*-phenylnaphthylamine (NPN, Sigma Aldrich) at a final concentration of 20 µM or propidium iodide (PI, Invitrogen, P3566) at a final concentration of 5 µM in darkness for 30 minutes at room temperature. Fluorescence was measured as follows: NPN Aex = 340 nm Aem = 420 nm and PI Aex = 535 nm Aem = 617 nm on a BioTek Synergy Neo2 Plate Reader. Relative fluorescence units (RFUs) were normalized to OD600.

#### Antibiotic uptake

To assess antibiotic uptake, the stationary phase cultures of the indicated cell lines in were diluted 1:50 into fresh LB Lennox media and cultured with diminazene at the indicated concentrations for 6 hours shaking at 37 °C. OD600 was measured and then cell lines were mixed with 50 µM of indicated antibiotic and cultured for 10 minutes shaking at 37 °C. Cells were immediately pelleted and then cell pellets were washed twice with PBS before being resuspended in LC–MS grade water (400 µL per 1 mL culture). Resuspended cells underwent two freeze thaw cycles alternating between liquid nitrogen and heating to 65 °C. Cell debris was pelleted and 90% of aqueous supernatant removed. LC–MS grade methanol (200 µL per 1 mL culture) was added to the cell debris. Cell debris was repelleted and methanol supernatant combined with the previously removed aqueous supernatant. This 2:1 water:methanol mix was centrifuged one additional time and the resulting supernatant used for LC–MS analysis (described below).

### *In vitro* biochemical experiments

#### Enzyme Expression and Purification

*SpeG* was amplified from *E. coli* K12 MG1655 genomic DNA and *SAT1*, *PA4114, PA1472,* and *PA1377* were ordered as *E. coli* codon optimized sequences (ThermoFisher); all were cloned cloned into pET-28A-inducible expression vectors using Gibson assembly (including an in-frame either N-terminal or C-terminal polyhistidine sequence) (primers in **Table S6**). Identities of the constructs were confirmed with DNA sequencing and then were transformed into *E. coli* BL21 (DE3) (New England Biolab) for expression. All *E. coli* expression constructs were grown overnight shaking at 37 °C in Terrific Broth (VWR) supplemented with 5 mM MgSO4 and kanamycin (50μg/ml) and were incubated at 37 °C overnight before dilution at 1:100 into fresh media the next morning. These diluted cultures were grown shaking at 37 °C to an OD600 of ∼0.6, at which point protein expression was induced by the addition of 200 µM isopropyl β-D-1-thiogalactopyranoside (IPTG, Teknova), followed by culturing shaking overnight at 16 °C. The next morning, *E. coli* were pelleted by centrifugation and then lysed in 20 mM HEPES pH 8.0 buffer containing 30 mM imidazole and 300 mM NaCl and supplemented with 0.5% octyl-β-D-thrioglucopyranoside (Chem-Impex), 0.5 mg/mL lysozyme (Sigma-Aldrich) and SIGFAST protease inhibitor cocktail (Sigma-Aldrich). After lysis and clarification by centrifugation, lysates were incubated for 1hour at 4 °C with His-Pure Cobalt Purification beads (Thermo Fisher Scientific). After incubation, beads were washed with six column volumes of 20 mM HEPES pH 8.0 buffer containing 30 mM imidazole and 300 mM NaCl before elution with one column volume of 20 mM HEPES pH 8.0 containing 300 mM imidazole and 300 mM NaCl. Eluted protein was then buffer exchanged into 20 mM HEPES pH 8.0 with 300 mM NaCl and 10% glycerol for storage at −80 °C before further experiments. Enzyme concentrations were calculated according to Beer’s Law with extinction coefficients calculated by Benchling software based on the amino acid sequences.

#### Enzyme assays

##### End point Assays

Assay mixtures contained 20 mM HEPES pH 7.5 with 50 mM NaCl with 1 µM enzyme, 50 mM putrescine, and 1 mM acetyl-CoA (CoALA Biosciences), supplemented with 1mM MgCl2 and 1mM Tris-(2- carboxyethyl)phosphine (TCEP, Sigma-Aldrich). Reactions were conducted at room temperature for 1 hour. Experiments were carried out in triplicate and repeated on different days with distinct protein preparations. Quenching/extraction was performed as below.

##### Determination of enzyme kinetics

For the determination of enzyme kinetics, a continuous spectrophotometric assay was used. Assay conditions were similar to those used in the above end point assays with 20 mM HEPES pH 7.5 with 50 mM NaCl supplemented with 1 mM MgCl2. Enzyme concentrations were varied based on the assay: 50 nM and 100 nM for SpeG with spermidine and putrescine respectively, 100 nM and 500 nM with SAT1 with spermidine and putrescine respectively, and 500 nM for PA1472 with putrescine. Concentrations of putrescine and spermidine were varied as indicated with 1 mM acetyl-CoA held constant. 100 µM 4,4’- dithiodipyridine (Sigma Aldrich) was added to measure the rate of coenzyme A generation (absorption maxima at 324 nm wavelength). Assays were carried out for 20-30 minutes at room temperature with absorption measured every 30-40 seconds. Each assay was performed in technical duplicates on the day of the experiment and results from distinct enzyme preparations tested on distinct days were averaged to produce the kinetic parameters.

<u>Determination of IC50 for diminazene.</u> For the determination of IC50s of diminazene for enzyme inhibition, the continuous enzyme assay from above was adapted with the modification that concentrations of substrate were held constant (near the empirically determined *K*halfs for SpeG, 50 mM putrescine and 600 µM spermidine, and *K*m for SAT1, 55 µM spermidine) and the concentration of diminazene was varied as indicated. As above, each assay was performed in technical duplicates on the day of the experiment and results from distinct enzyme preparations tested on distinct days were averaged to produce the kinetic parameters.

#### Mass spectrometry

##### Putrescine metabolites

Plasma samples, (spent) media, and end-point enzyme assays were quenched/extracted with one part sample and nine parts extraction mix (74.9% acetonitrile: 24.9% methanol: 0.2% formic acid v/v/v with 2 µg/mL of valine-d8 and phenylalanine-d8 from Cambridge Isotopes as an internal standard for quantification). For intracellular putrescine accumulation, cells were extracted via multiple freeze thaw cycles with water and methanol (final 67% water: 33% methanol v/v). Extracted/quenched samples were vortexed and cooled to −20 °C and then centrifuged before LC–MS analysis. MS analyses were conducted using an LC–MS system composed of an Agilent 1290 Infinity II UHPLC (capable of column switching) coupled to an Agilent 6470A Triple Quadrupole LC/MS. The samples were injected into an Infinity Lab Poroshell 120 HILIC column (2.1×100mm×2.7µm) at 25 °C. The column was eluted isocratically at a flow rate of 600 µL min^−1^ with 5% mobile phase A (10mM ammonium formate with 0.1% formic acid in water) for 18 seconds, followed by a linear gradient for 132 seconds to 60% mobile phase B (acetonitrile with 0.1% formic acid). This was followed by a 3-second gradient to 40% mobile phase B. This was then followed by a 27-second gradient returning to 5% mobile phase A at a flow rate of 1,200 µL/min. This flow rate and ratio was held for an additional 48seconds. Additional column equilibration was carried out on a secondary pump for 117 seconds at 5% mobile phase A and a flow rate of 1,000 µL/min. MS was conducted in positive ion mode using ESI. Data were collected via MRM (**Table S7**). Other MS settings were as follows: heater temp 300 °C, ESI nebulizer 45 psi, spray voltage 3.5 kV and acquisition time 75 ms per spectrum. Raw data from the LC–MS were analyzed using Agilent MassHunter Quantitative Analysis version 10.1 software. For absolute quantification, all samples were normalized to the valine and phenylalanine internal standards and concentrations calculated against a standard curve. Standard curves were generated using putrescine dihydrochloride and *N*-acetylputrescine hydrochloride purchased from Sigma Aldrich, and lysophastidylcholine 16:0 and 18:0 purchased from Avanti Polar Lipids. For all cell culture experiments, metabolite measurements were further normalized to OD600.

##### Antibiotic uptake

MS analyses were conducted using an LC–MS system composed of an Agilent 1290 Infinity II UHPLC (capable of column switching) coupled to an Agilent 6470A Triple Quadrupole LC/MS. The samples were injected into an Infinity Lab Poroshell 120 C18 column (2.1×50mm×2.7µm) at 30 °C. The column was eluted isocratically at a flow rate of 200 µL/min with 90% mobile phase A (0.1% formic acid in water) for 1 minute, followed by a linear gradient for 4 minutes to 98% mobile phase B (acetonitrile with 0.1% formic acid). This was followed by 1 minute of isocratic elution at 98% mobile phase B, before a 6 second gradient returning to 90% mobile phase A at a new flow rate of 400 µL/min. The column was then eluted isocratically with 90% mobile phase A for 1.4 minutes at a flow rate of 400 µL/min. MS was conducted in positive ion mode using ESI. Data were collected via MRM (**Table S7**). Other MS settings were as follows: heater temp 300 °C, ESI nebulizer 45 psi, spray voltage 3.5 kV and acquisition time 200 ms per spectrum. For all cell culture experiments, measurements were normalized to OD600.

### Computational Experiments

#### Identification of PA1472 and PA1377

The *E. coli* SpeG and *B. subtilis* PaiA and BltD protein sequences were used in a Basic Local Alignment Search Tool (BLAST) search of the Joint Genome Institute – Integrated Microbial Genomes database^109^ of *P. aeruginosa* PAO1 isolates. SpeG as a query identified PA1472 and PA1377 with alignment scores of 7e-10 and 3e-9 respectively. PaiA as a query identified no hits. BltD identified PA4114 and PA1377 with alignment scores of 2e-16 and 4e-8 respectively.

#### Phylogenetic tree

All sequences from Uniprot (http://uniprot.org) tagged with EC 2.3.1.57 and spermidine or spermidine were downloaded in FASTA format. Analysis was limited to sequences with between 140 and 200 amino acids. A multiple sequence alignment was generated using MUSCLE^49^ v3.8.425 on Geneious Prime 2022.2.1 (Dotmatics) of the 1483 representatives of enzyme groups sharing >80% amino acid sequence identity in the UniProt database (release 2023_2) using EC: 2.3.1.57 AND “spermidine or spermidine” as a query. PA1472 and PA1377 were also included in this analysis. A maximum-likelihood phylogenetic tree was constructed using iQ-TREE2 v2.1.0 with model finder^51,110^, which determined VT+R10 as the best model, and visualized using iTOL^50^. Branch supports were calculated from 1001 independent tree iterations.

#### Protein structures

AlphaFold2^111^ was used to generate a predicted homodimeric structures for PA1472 using sequence Q9I3P0_PSEAE downloaded from the UniProt database. Published sequences of *E. coli* SpeG (3WR7) and *H. sapiens* SAT1 (2B5G) were downloaded from the Protein Data Bank (PBD) and used for comparison^44,53^. Alignments were conducted using the dimer structures of each protein and “align” tool in Pymol 2.5.3.

#### Statistical Analyses

Separate from our human analyses, appropriate statistical tests were performed where indicated. All analyses were carried out using GraphPad Prism 9 (GraphPad Software).

---

**Figure S1.**
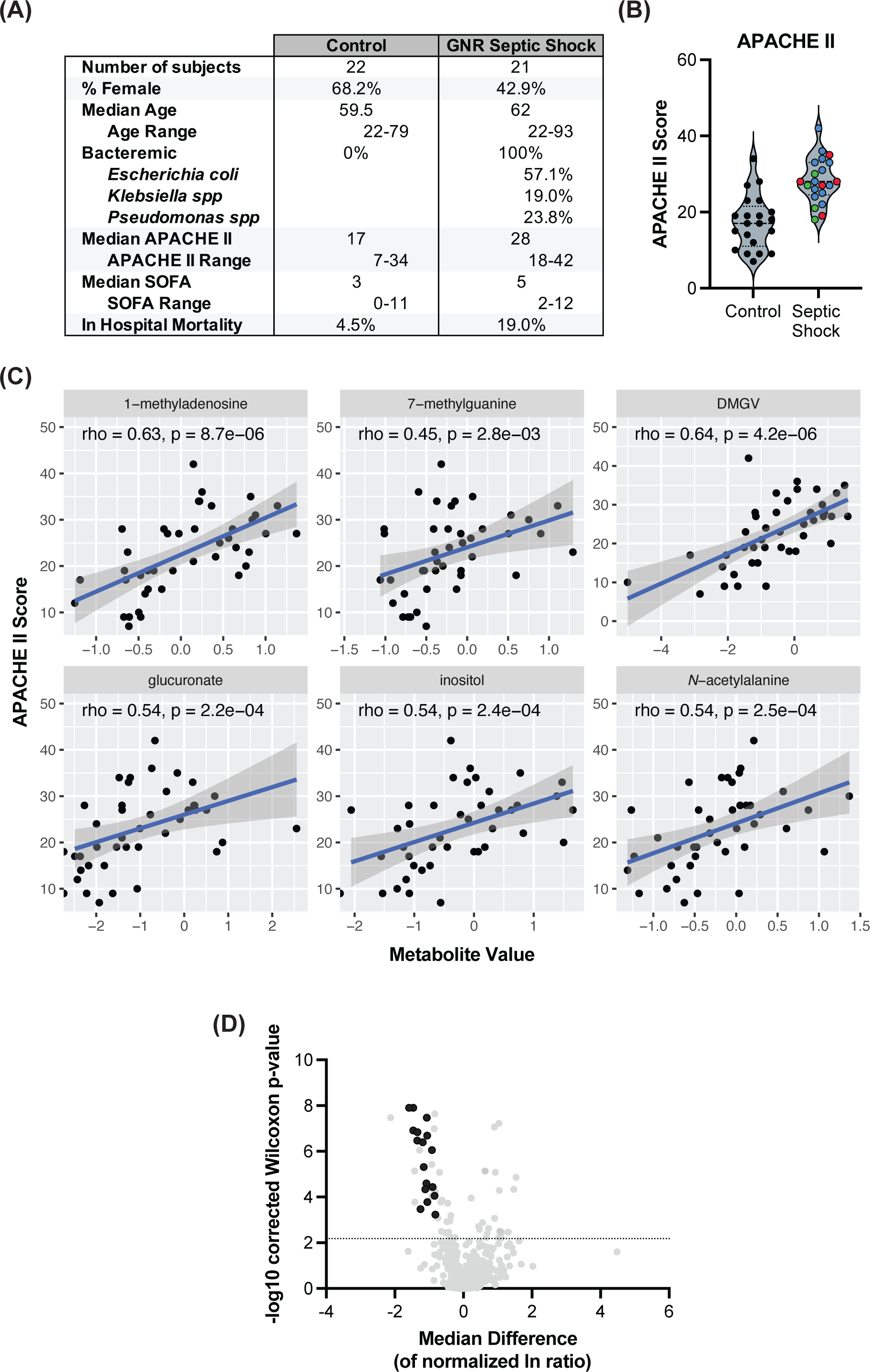
Putrescine metabolites are elevated in humans with gram-negative septic shock. (A) Patient characteristics (B) APACHE II distribution by organism; Blue = *E. coli*, Green = *K. pneumoniae*, Red = *P. aeruginosa* (C) Correlation plots for the other 6 metabolites of significance that also showed significant correlations with APACHE II; DMGV = dimethylguanidino valerate (D) Volcano plot with highlighting significant decreases in numerous lysophosphatidylcholine species

**Figure S2.**
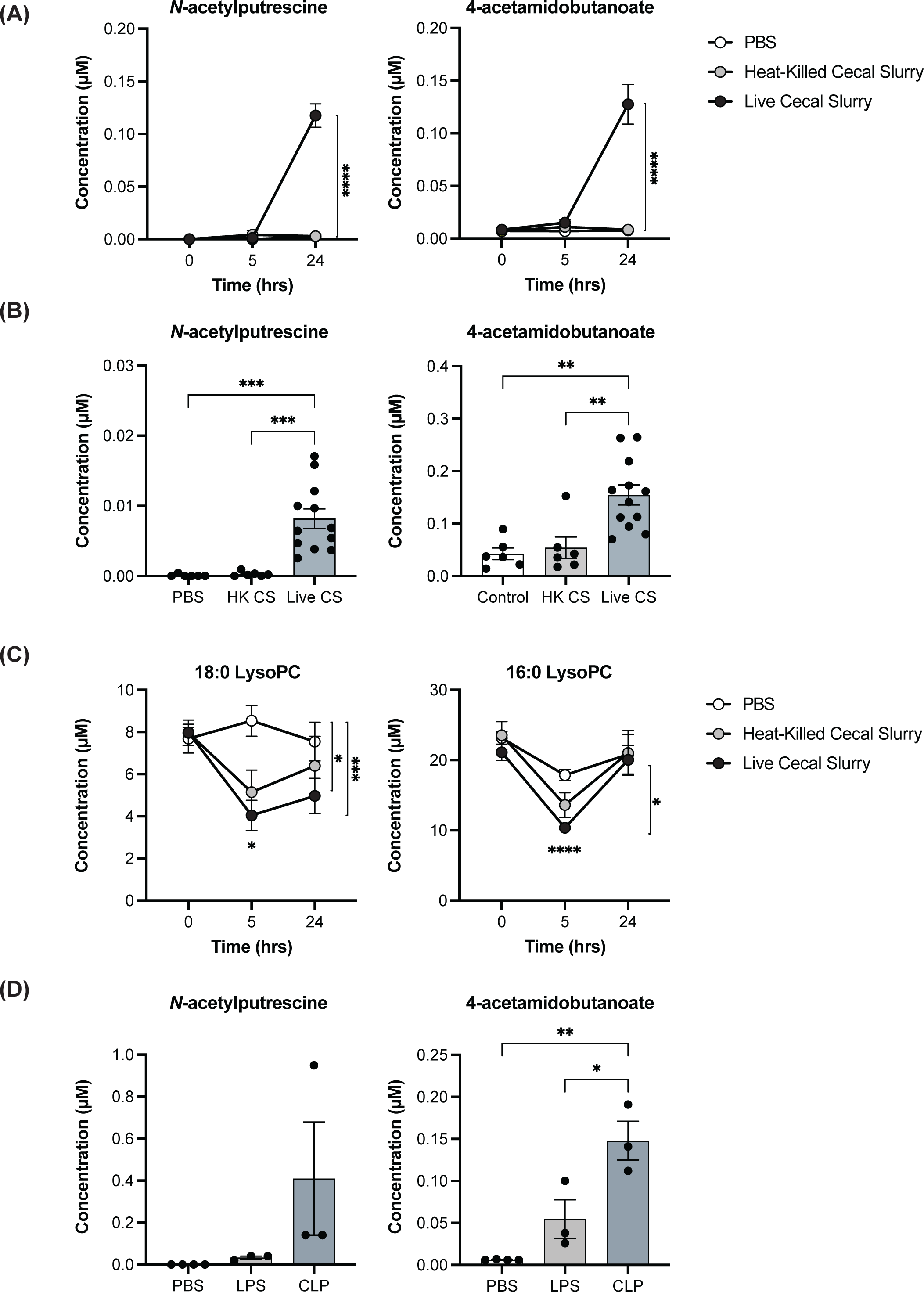
Putrescine metabolites are elevated in mouse models of septic shock. (A) Time course of putrescine metabolite changes in cecal slurry model; HK = heat killed, CS = cecal slurry; n= 8 PBS, n = 7 HK CS, n = 4 Live CS; p-values were determined by Two-way ANOVA followed by Tukey’s multiple comparisons test (B) Putrescine metabolites are elevated in the mouse cecal slurry model of septic shock in female mice; HK = heat killed, CS = cecal slurry; n = 6 PBS, n = 6 HK CS, n = 12 Live CS; p-values were determined by One-way ANOVA followed by Tukey’s multiple comparisons test (C) Time course of representative lysophosphatidylcholine (lysoPC) changes in cecal slurry model; HK = heat killed, CS = cecal slurry; n = 8 PBS, n = 7 HK CS, n = 4 Live CS; p-values were determined by Two-way ANOVA followed by Tukey’s multiple comparisons test (D) Cecal ligation and puncture 24 hour time point demonstrates similar elevations in putrescine metabolites; n = 4 PBS, n = 3 lipopolysaccaride (LPS), n = 3 cecal ligation and puncture (CLP). p-values were determined by One-way ANOVA followed by Tukey’s multiple comparisons test (E) For all panels, data presented are means ± SEM; *p < 0.05; **p < 0.01; ***p < 0.001; ****p<0.0001

**Figure S3.**
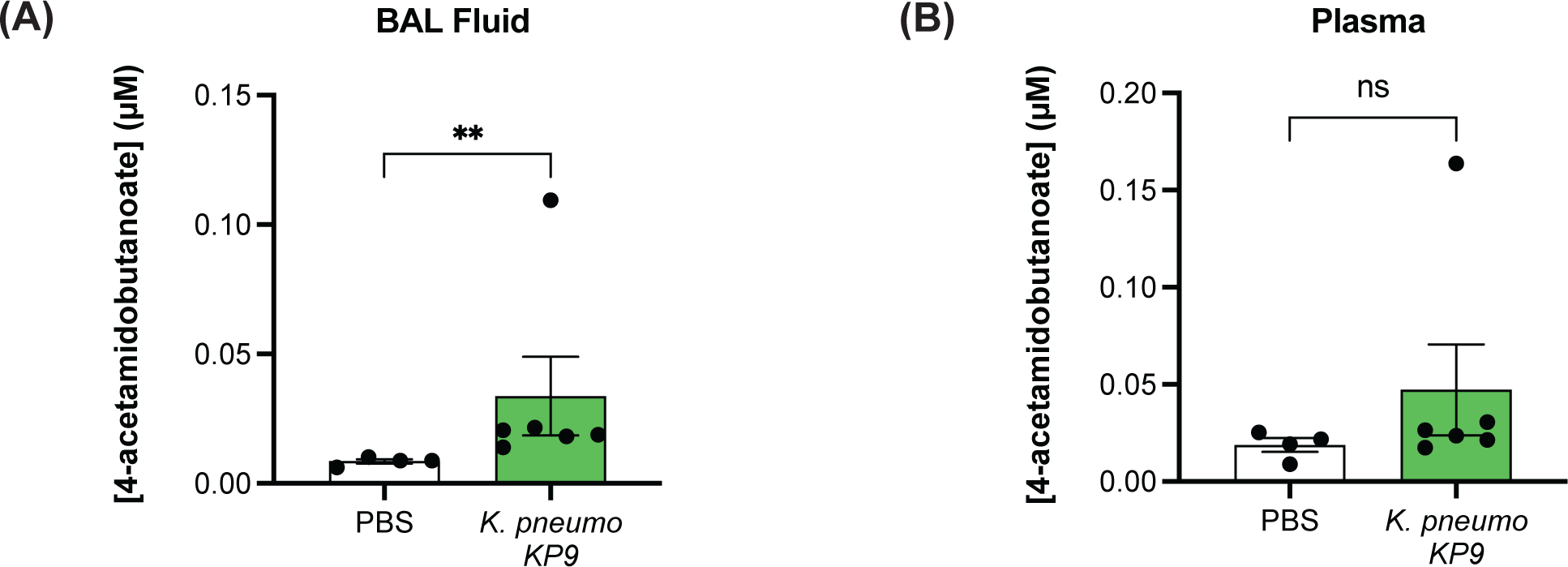
*Klebsiela pneumoniae* pneumonia levels of 4-acetamidobutanoate. (A) *Klebsiella pneumoniae* pneumonia results in increased bronchial alveolar lavage (BAL) fluid levels of 4- acetamidobutanoate; n = 4 PBS, n = 6 *K. pneumoniae* (KP9); Two-tailed p-values were determined by Mann Whitney test to include outlier. (B *Klebsiella pneumoniae* pneumonia results in trend towards increased plasma levels of 4- acetamidobutanoate; n = 4 PBS, n = 6 *K. pneumoniae* (KP9); Two-tailed p-values were determined by Mann Whitney test to include outlier. For all panels, data presented are means ± SEM

**Figure S4.**
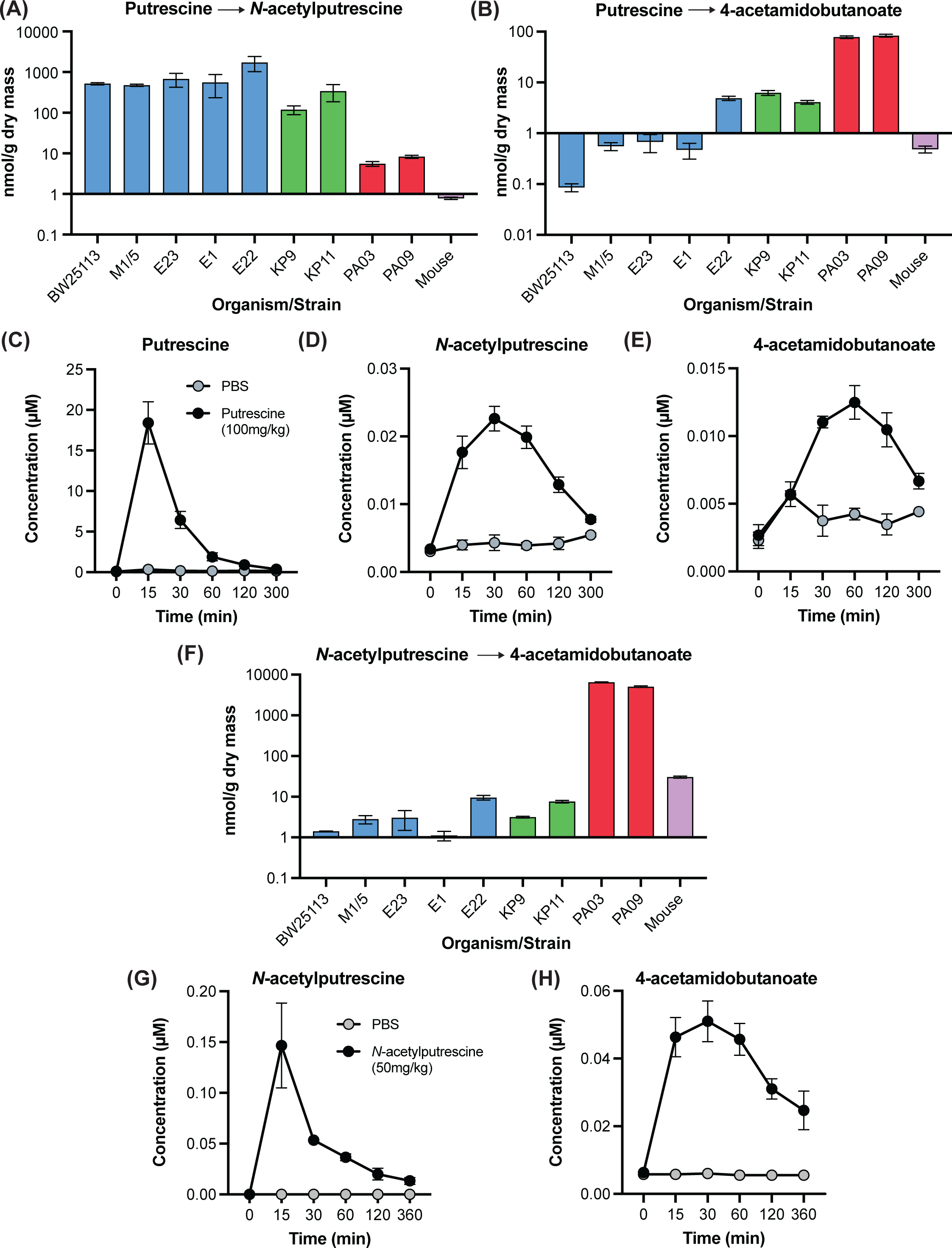
Putrescine metabolite production by bacteria and mice. (A) Production of *N*-acetylputrescine from putrescine by bacteria and mice; n = 3 for bacteria, representative data from 1-3 independent repeats; n = 6 mice (B) Production of 4-acetamidobutanoate from putrescine by bacteria and mice; n = 3 for bacteria, representative data from 1-3 independent repeats; n = 6 mice (C) Time course of putrescine levels in mouse plasma after IP injection; n = 6 mice for each treatment (D) Time course of *N*-acetylputrescine plasma levels in mice after IP injection with 100 mg/kg of putrescine; n = 6 mice for each treatment (E) Time course of 4-acetamidobutanoate plasma levels in mice after IP injection with 100 mg/kg of putrescine; n = 6 mice for each treatment (F) Production of 4-acetamidobutanoate from *N*-acetylputrescine by bacteria and mice; n = 3 for bacteria, representative data from 1-3 independent repeats; n = 3 mice (G) Time course of *N*-acetylputrescine plasma levels in mice after IP injection with 50 mg/kg of *N*- acetylputrescine; n = 4 PBS, n = 3 *N*-acetylputrescine (H) Time course of 4-acetamidobutanoate plasma levels in mice after IP injection with 50 mg/kg of *N*- acetylputrescine; n = 4 PBS, n = 3 *N*-acetylputrescine For all panels, data presented are means ± SEM

**Figure S5.**
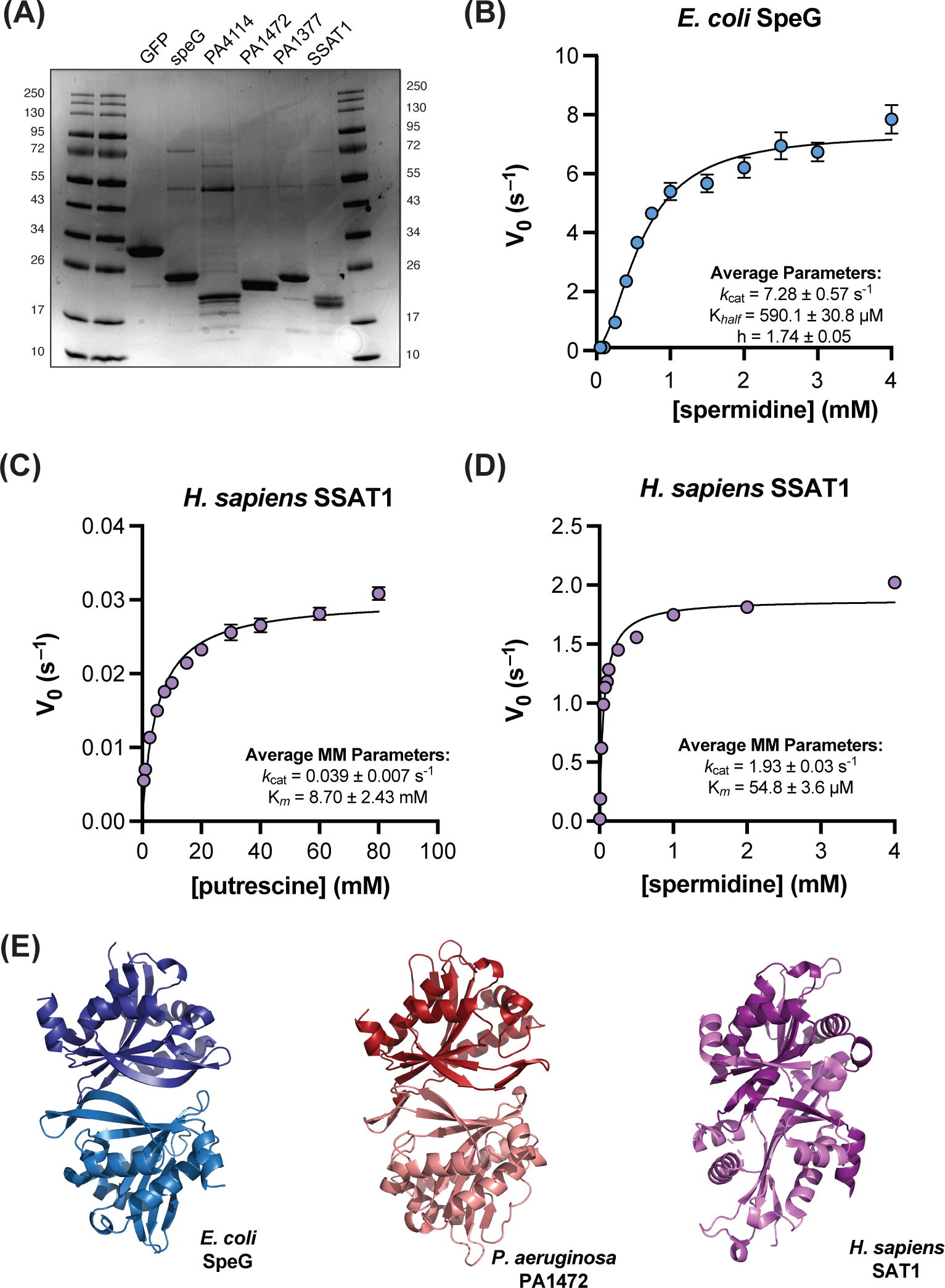
SpeG homologs are responsible for production of *N*-acetylputrescine in bacteria and are distinct from mammals. (A) Uncropped protein gel of purified proteins (B) Kinetics of SpeG on spermidine; n = 3 per concentration, representative data from 3 independent experiments; summary parameters includes all experiments (C) Kinetics of SAT1 on putrescine; n = 3 per concentration, representative data from 4 independent experiments; summary parameters includes all experiments (D) Kinetics of SAT1 on spermidine; n = 3 per concentration, representative data from 3 independent experiments; summary parameters includes all experiments (E) SpeG (PDB: 3WR7), PA1472 (AlphaFold2), SAT1 (PDB: 2B5G) dimers For all panels, data presented are means ± SEM

**Figures S6.**
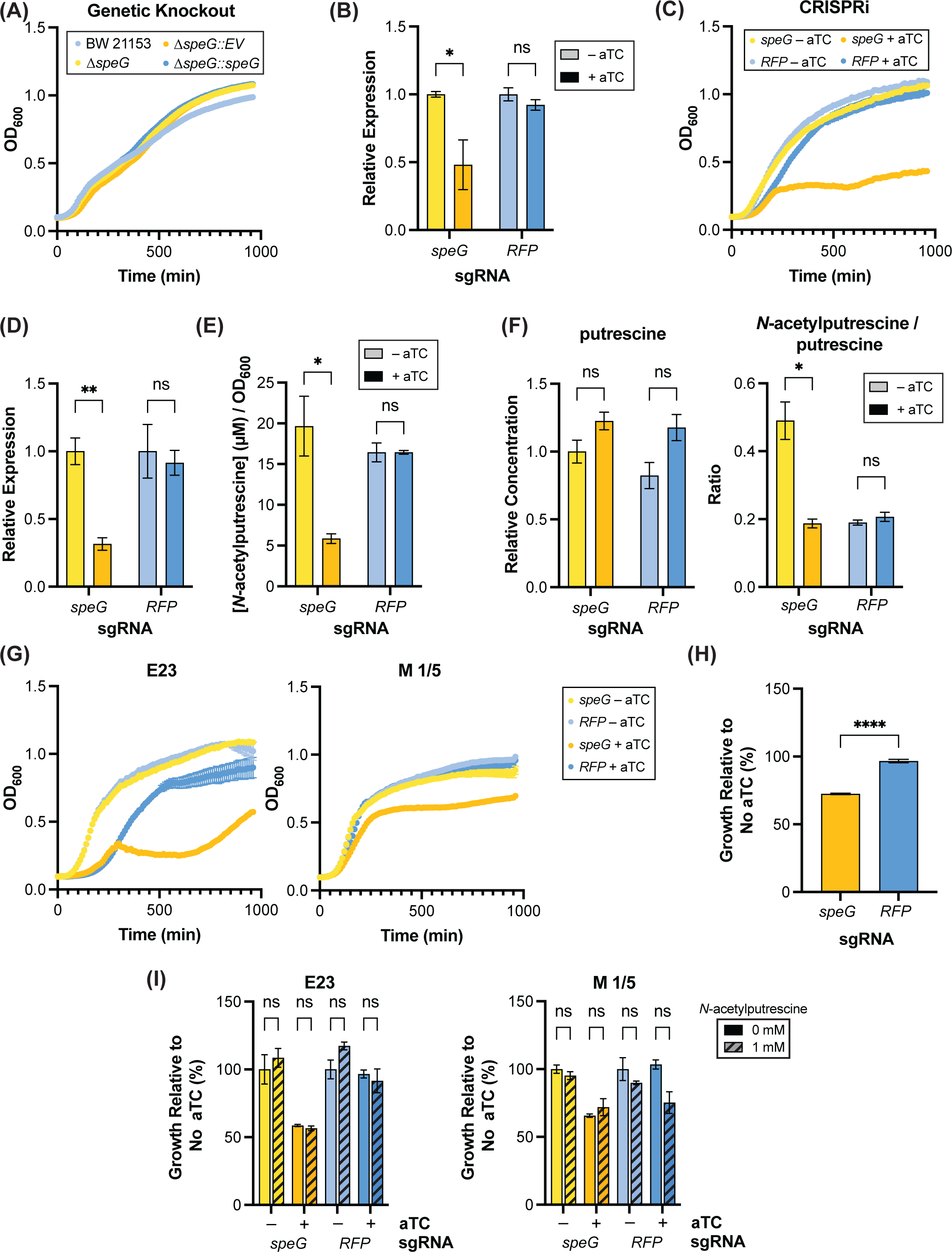
Suppresion of SpeG expression impacts proliferation. (A) *E. coli* BW25113 parental strain growth compared with KEIO collection *ΔspeG* suggests no clear effect on proliferation; n = 3 per cell line; representative data from 3 independent experiments (B) Suppression of *speG* expression with inducible CRISPRi in BW25113; n = 3 per condition, representative data from 2 independent experiments (C) Inducible CRISPRi of *speG* in BW25113 demonstrates suppression of *speG* expression impairs proliferation; n = 3 per condition, representative data from 2 independent experiments (D) Suppression of *speG* expression with inducible CRISPRi in M 1/5; n = 3 per condition, representative data from 3 independent experiments (E) Decreased levels of extracellular *N*-acetylputrescine with inducible CRISPRi of *speG* in M 1/5, concentrations normalized to OD600 = 1.0; n = 3 per condition, representative data from 2 independent experiments (F) Inducible CRISPRi of *speG* blocks putrescine to *N*-acetylputrescine conversion intracellularly in M 1/5; n = 3 per condition (G) Raw growth curves for inducible CRISPRi in M 1/5 and E23 (Figure 3D and Figure S6H) (H) Inducible CRISPRi of *speG* suppress growth in *E. coli* isolate M 1/5; n = 3 per condition, representative data from 2 independent experiments (I) Supplementation of inducible CRISPRi cultures of *speG* with 1mM *N*-acetylputrescine does not rescue proliferation defect; n = 3 per condition, representative data from 3 independent experiments For all panels data presented are means ± SEM; Two-tailed p-values were determined by unpaired t test; *p < 0.05; **p < 0.01; ***p < 0.001; ****p<0.0001 aTC = anhydrotetracycline

**Figure S7.**
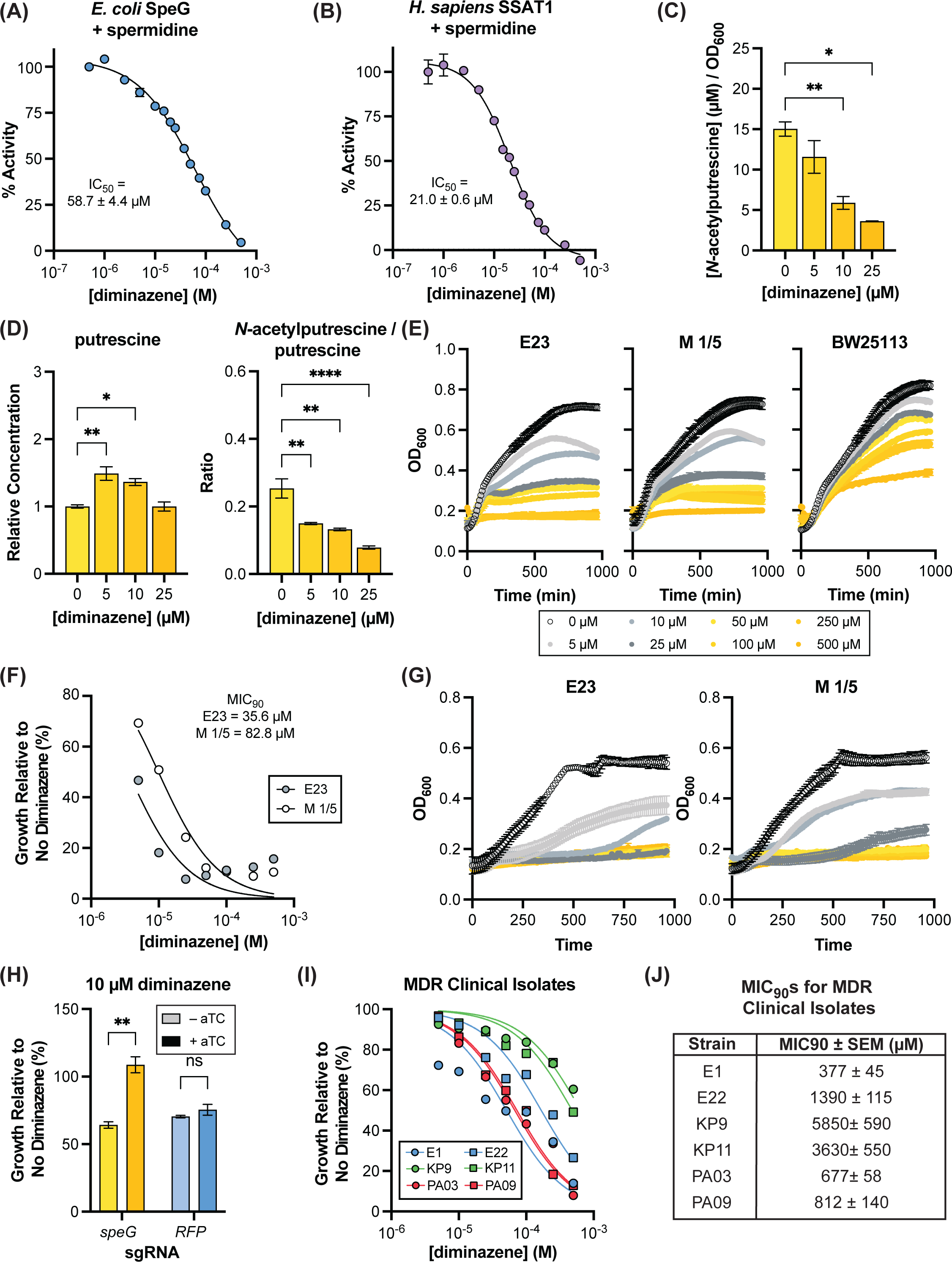
Diminazene inhibits SpeG and phenocopies inducible CRISPRi. (A) Determination of *in vitro* IC50 of diminazene against SpeG with spermidine substrate; n = 2 per inhibitor concentration, representative data from 3 independent experiments; summary parameter includes all experiments (B) Determination of *in vitro* IC50 of diminazene against SAT1 with spermidine substrate; n = 2 per inhibitor concentration, representative data from 3 independent experiments; summary parameter includes all experiments (C) Diminazene treatment reduces extracellular levels of *N*-acetylputrescine in M 1/5, concentrations normalized to OD600 = 1.0; n = 3 per condition, representative data from 3 independent experiments; p- values were determined by One-way ANOVA followed by Dunnett’s multiple comparisons test with all comparisons made against no drug control (D) Diminazene treatment increases relative intracellular putrescine levels and decreases the ratio of *N*- acetylputrescine in M 1/5; n = 3 per condition; normalized to OD600 = 1.0, then normalized to the concentration of the control, 0 µM, condition; p-values were determined by One-way ANOVA followed by Dunnett’s multiple comparisons test with all comparisons made against no drug control (E) Growth curves for diminazene treatment MIC90s of E23, M 1/5, and BW25113 in LB; n = 3 per condition (F) MIC90 of E23 and M 1/5 in M9 minimal media with 0.4% glucose and 0.2% Cas-AA; n = 3 per condition, representative data from 2 independent experiments; summary MIC90s combine all experiments (G) Representative growth curves for diminazene treatment MIC90s of E23 and M 1/5 in M9 minimal media with 0.4% glucose and 0.2% Cas-AA (Figure S7F); n = 3 per condition (H) Inducible CRISPRi of *speG* blocks growth inhibition by diminazene in M 1/5; n = 3 per condition, presentative data from 3 independent experiments (I) MIC curves of MDR clinical isolates of *E. coli, K. pneumonia*, *P. aeruginosa* in LB; n = 3 per condition, representative data from 3 independent experiments Calculated MIC90 values for MDR patient isolates in LB (Figure S7I) For all panels data presented are means ± SEM; *p < 0.05; **p < 0.01; ***p < 0.001; ****p<0.0001 aTC = anhydrotetracycline

**Figure S8.**
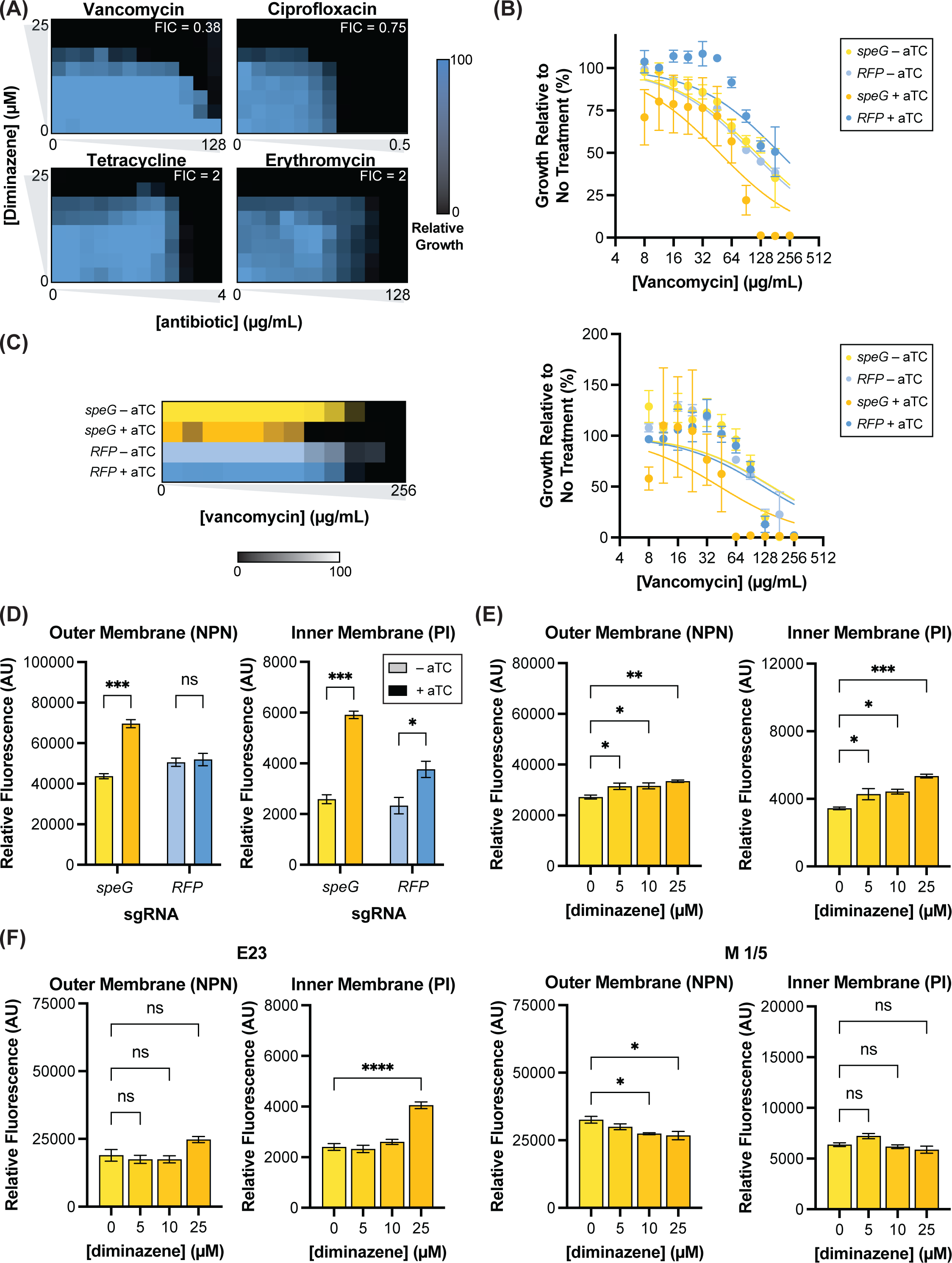
Reducing SpeG activity enhances membrane permeability. (A) Checkerboard assays demonstrate synergy between diminazene and vancomycin in M 1/5; representative data from 4 independent experiments (B) Growth relative to no vancomycin of inducible CRISPRi of *speG* in E23 in LB with vancomycin gradient; n = 3 per condition, representative data from 3 independent experiments (graphical summary of data in Figure 5B). (C) Inducible CRISPRi of *speG* reduces MIC of vancomycin in M 1/5 and corresponding growth curves relative to no vancomycin; mean of n = 3 per condition, representative data from 3 independent experiments (D) Inducible CRISPRi of *speG* enhances membrane permeability of M 1/5; n = 3 per condition, representative data from 2 independent experiments; Two-tailed p-values were determined by unpaired t test (E) Diminazene treatment for 6 hours enhances membrane permeability in M1/5; n = 3 per condition, representative data from 2 independent experiments; p-values were determined by One-way ANOVA followed by Dunnett’s multiple comparisons test with all comparisons made against no drug control (F) Diminazene treatment for 3 hours has minimal to no effect on membrane permeability in E23 and M 1/5; n = 3 per condition; p-values were determined by One-way ANOVA followed by Dunnett’s multiple comparisons test with all comparisons made against no drug control For B-F, data presented are means ± SEM; *p < 0.05; **p < 0.01; ***p < 0.001; ****p<0.0001 aTC = anhydrotetracycline

**Figure S9.**
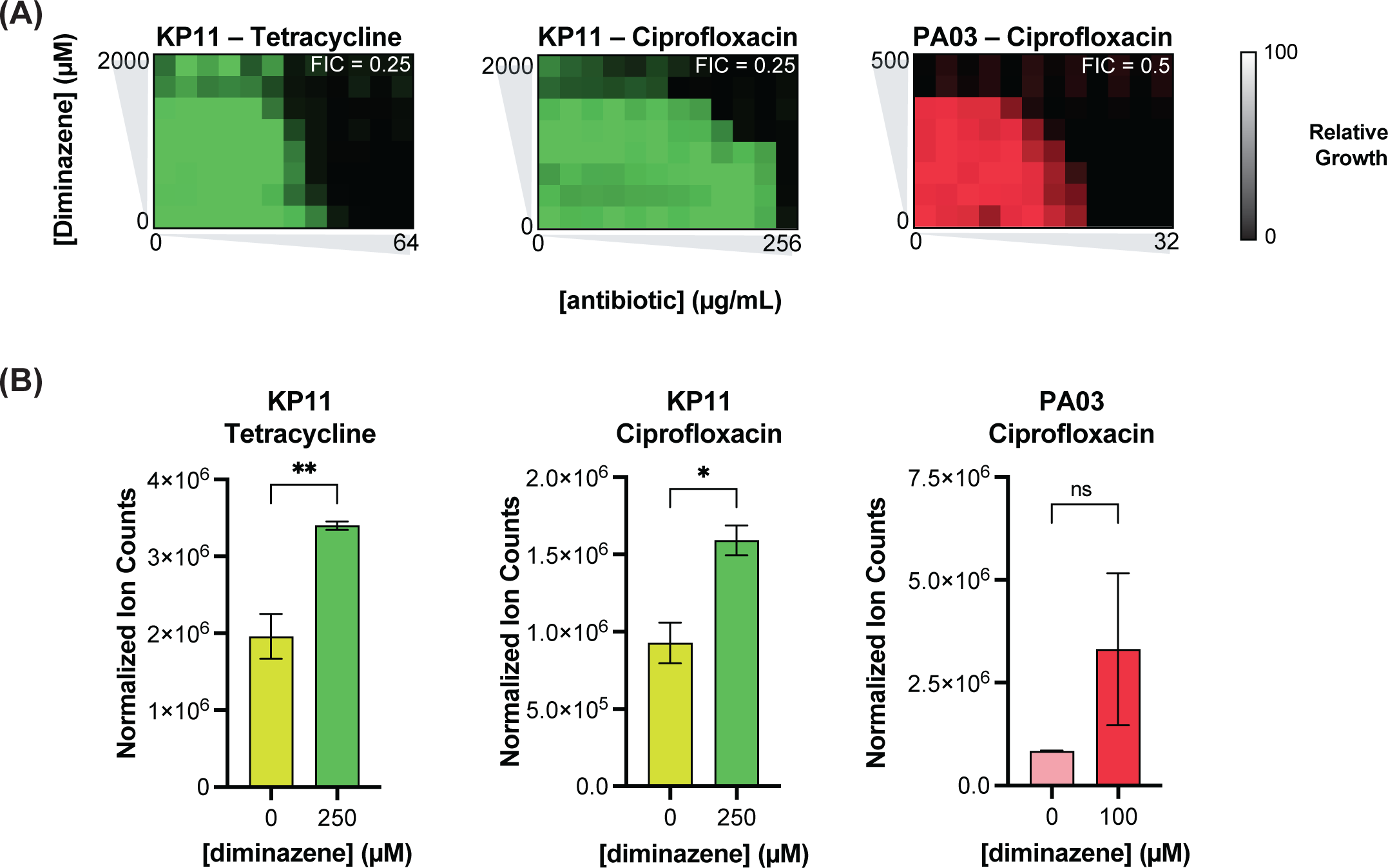
Blocking SpeG synergizes with existing clinical antibiotics in resistant bacteria. (A) Checkerboard assays demonstrate synergy between diminazene and antibiotics to which the assayed MDR strains are clinically resistant; representative data from 1-3 independent experiments per cell line (B) Diminazene treatment enhances uptake of antibiotics to which MDR strains are resistant; n = 3 per condition, representative data from 2 independent experiments; Two-tailed p-values were determined by unpaired t test For panel B, data presented are means ± SEM; *p < 0.05; **p < 0.01; ***p < 0.001; ****p<0.0001 aTC = anhydrotetracycline

**Table S1.** Individual patient characteristics.

| Subject ID | Age | Sex | Group | Diagnosis | Microorganism | Baseline APACHE II | Baseline SOFA | In hosp mortality? |
| --- | --- | --- | --- | --- | --- | --- | --- | --- |
| 1 | 64 | F | Septic Shock | Septic shock | *Escherichia coli* | 28 | 6 | N |
| 2 | 93 | M | Septic Shock | Septic shock | *Pseudomonas species* | 28 | 4 | N |
| 3 | 87 | F | Septic Shock | Septic shock | *Klebsiella species* | 30 | 4 | N |
| 4 | 44 | F | Septic Shock | Septic shock | *Pseudomonas species* | 28 | 3 | Y |
| 5 | 53 | M | Septic Shock | Septic shock | *Escherichia coli* | 24 | 2 | N |
| 6 | 60 | M | Septic Shock | Septic shock | *Klebsiella species* | 18 | 5 | Y |
| 7 | 26 | M | Control | Crohn's, desensitization |  | 9 | 0 | N |
| 8 | 73 | M | Septic Shock | Septic shock | *Escherichia coli* | 36 | 7 | N |
| 9 | 72 | M | Septic Shock | Septic shock | *Escherichia coli* | 26 | 2 | N |
| 10 | 68 | F | Control | Aspirin desensitization |  | 9 | 0 | N |
| 11 | 62 | M | Septic Shock | Septic shock | *Escherichia coli* | 42 | 9 | Y |
| 12 | 56 | F | Septic Shock | Septic shock | *Pseudomonas species* | 27 | 2 | N |
| 13 | 45 | M | Septic Shock | Septic shock | *Pseudomonas species* | 35 | 9 | N |
| 14 | 85 | F | Septic Shock | Septic shock | *Escherichia coli* | 33 | 6 | N |
| 15 | 58 | M | Septic Shock | Septic shock | *Pseudomonas species* | 19 | 3 | N |
| 16 | 42 | F | Septic Shock | Septic shock | *Klebsiella species* | 27 | 4 | N |
| 17 | 57 | M | Septic Shock | Septic shock | *Escherichia coli* | 27 | 12 | N |
| 18 | 58 | F | Septic Shock | Septic shock | *Escherichia coli* | 31 | 4 | Y |
| 19 | 43 | M | Control | Volume overload, possible pneumonia |  | 34 | 8 | N |
| 20 | 22 | F | Septic Shock | Septic shock | *Klebsiella species* | 21 | 4 | N |
| 21 | 84 | F | Septic Shock | Septic shock | *Escherichia coli* | 34 | 9 | N |
| 22 | 58 | F | Control | Angioedema |  | 15 | 3 | N |
| 23 | 61 | F | Control | ALS |  | 14 | 3 | N |
| 24 | 57 | M | Control | Hyponatremia and alcohol intoxication |  | 23 | 7 | N |
| 25 | 64 | F | Control | Upper GI bleed |  | 10 | 5 | N |
| 26 | 68 | F | Control | Bleeding after interventional radiology procedure |  | 20 | 3 | N |
| 27 | 71 | M | Septic Shock | Septic shock | *Escherichia coli* | 25 | 6 | N |
| 28 | 89 | M | Septic Shock | Septic shock | *Escherichia coli* | 33 | 6 | N |
| 29 | 79 | M | Control | Mouth bleeding |  | 15 | 2 | N |
| 30 | 58 | M | Control | Status post cardiac arrest |  | 7 | 3 | N |
| 31 | 57 | F | Control | Respiratory failure from influenza (cultures negative) |  | 17 | 1 | N |
| 32 | 66 | F | Control | COPD exacerbation |  | 27 | 4 | N |
| 33 | 57 | M | Control | Pneumonitis |  | 28 | 11 | Y |
| 34 | 42 | F | Control | Hyponatremia with PCP pneumonia |  | 19 | 5 | N |
| 35 | 75 | F | Control | Tumor impinging on trachea |  | 18 | 0 | N |
| 36 | 68 | F | Control | RF and ILD admitted for lung transplant evaluation |  | 17 | 2 | N |
| 37 | 22 | F | Control | Asthma exacerbation |  | 12 | 0 | N |
| 38 | 68 | F | Control | Respiratory failure from SCLC |  | 19 | 0 | N |
| 39 | 91 | M | Septic Shock | Septic shock | *Escherichia coli* | 22 | 9 | N |
| 40 | 30 | F | Control | Influenza B complicated by ARDS |  | 23 | 10 | N |
| 41 | 31 | F | Control | Pancreatitis from hypertriglyceridemia |  | 9 | 2 | N |
| 42 | 76 | M | Control | Community acquired pneumonia, rule out MERS |  | 19 | 3 | N |
| 43 | 61 | F | Control | Pulmonary hypertension admitted for lung transplant evaluation |  | not available | not available | N |

**Table S2.**
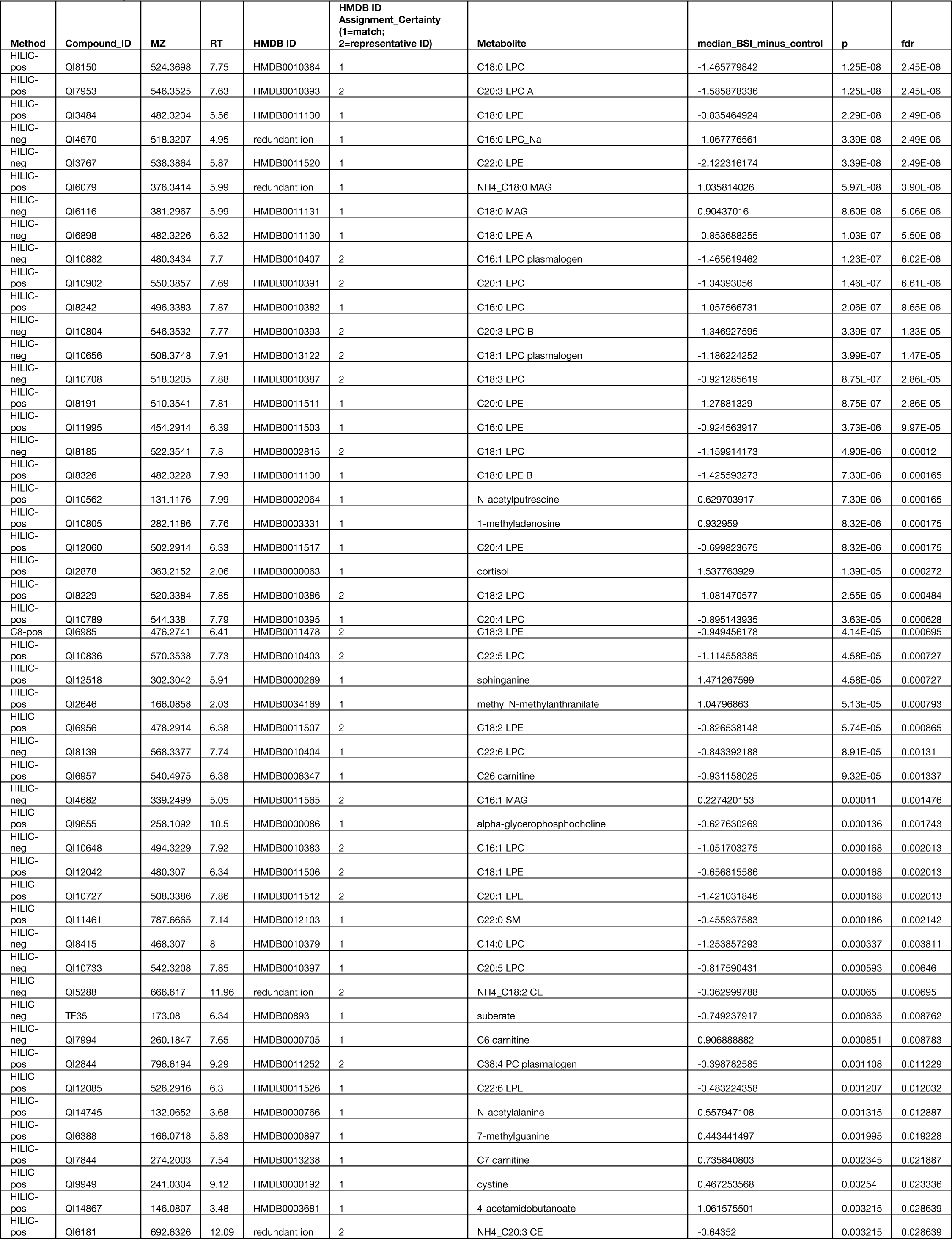
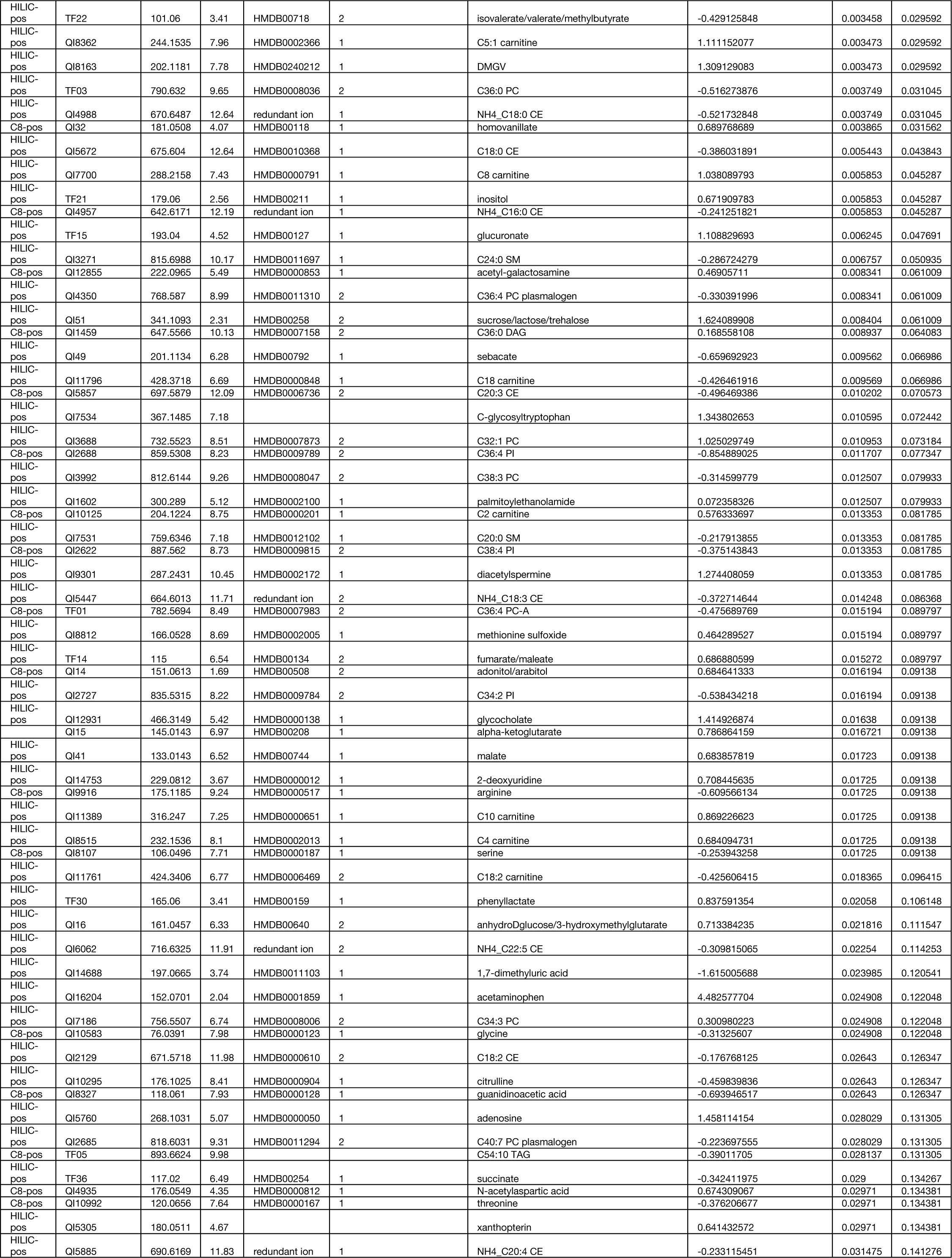
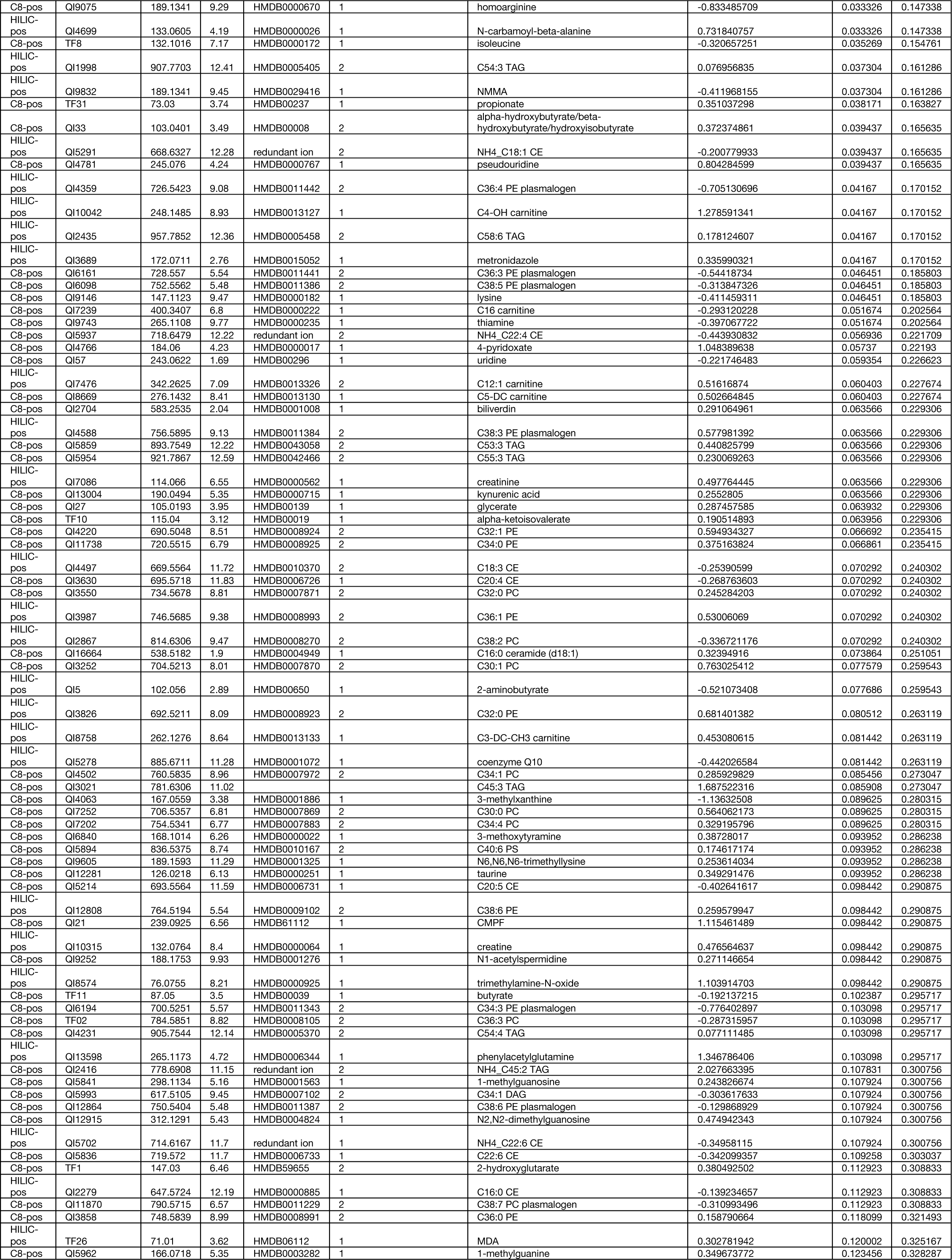
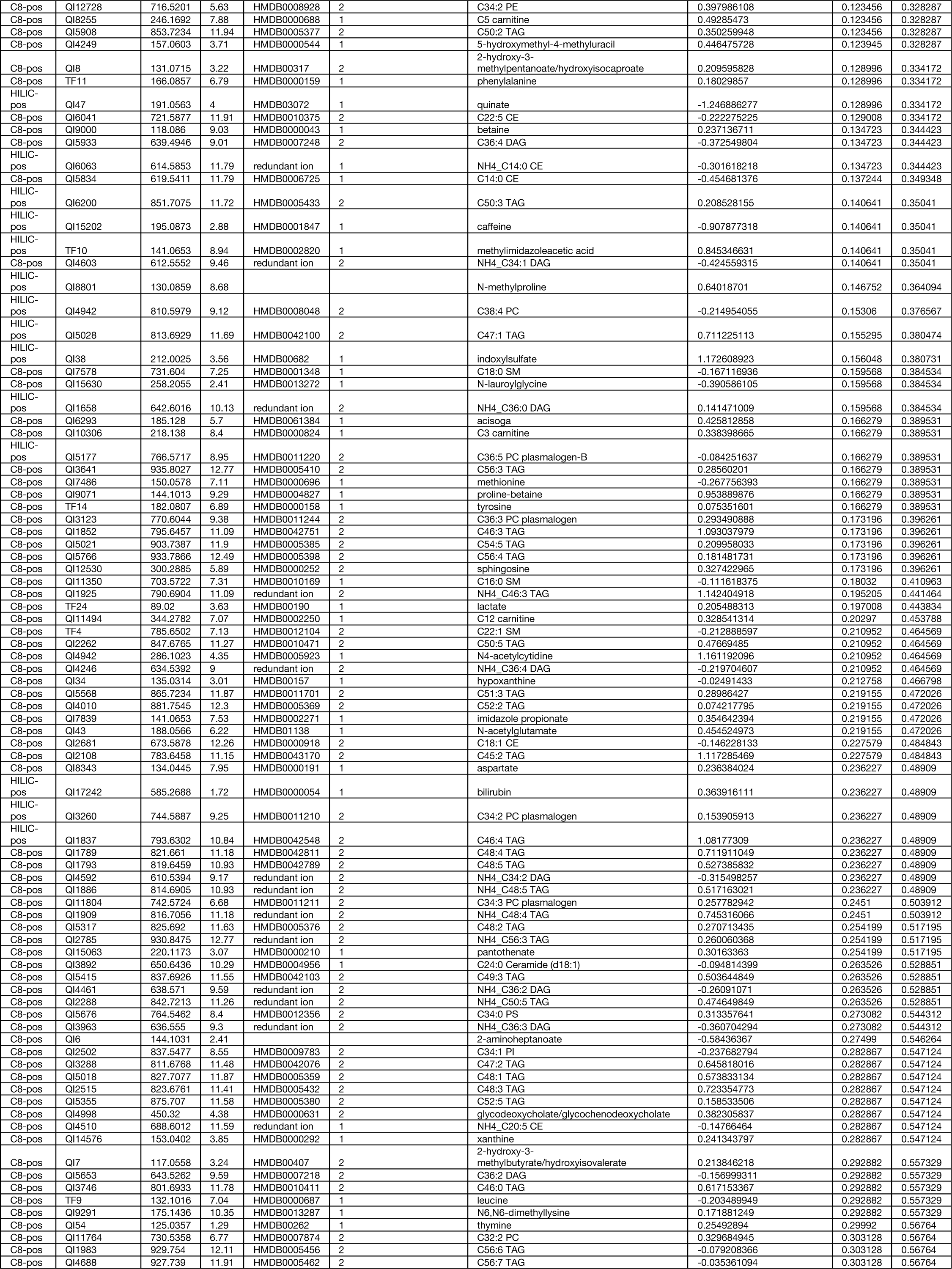
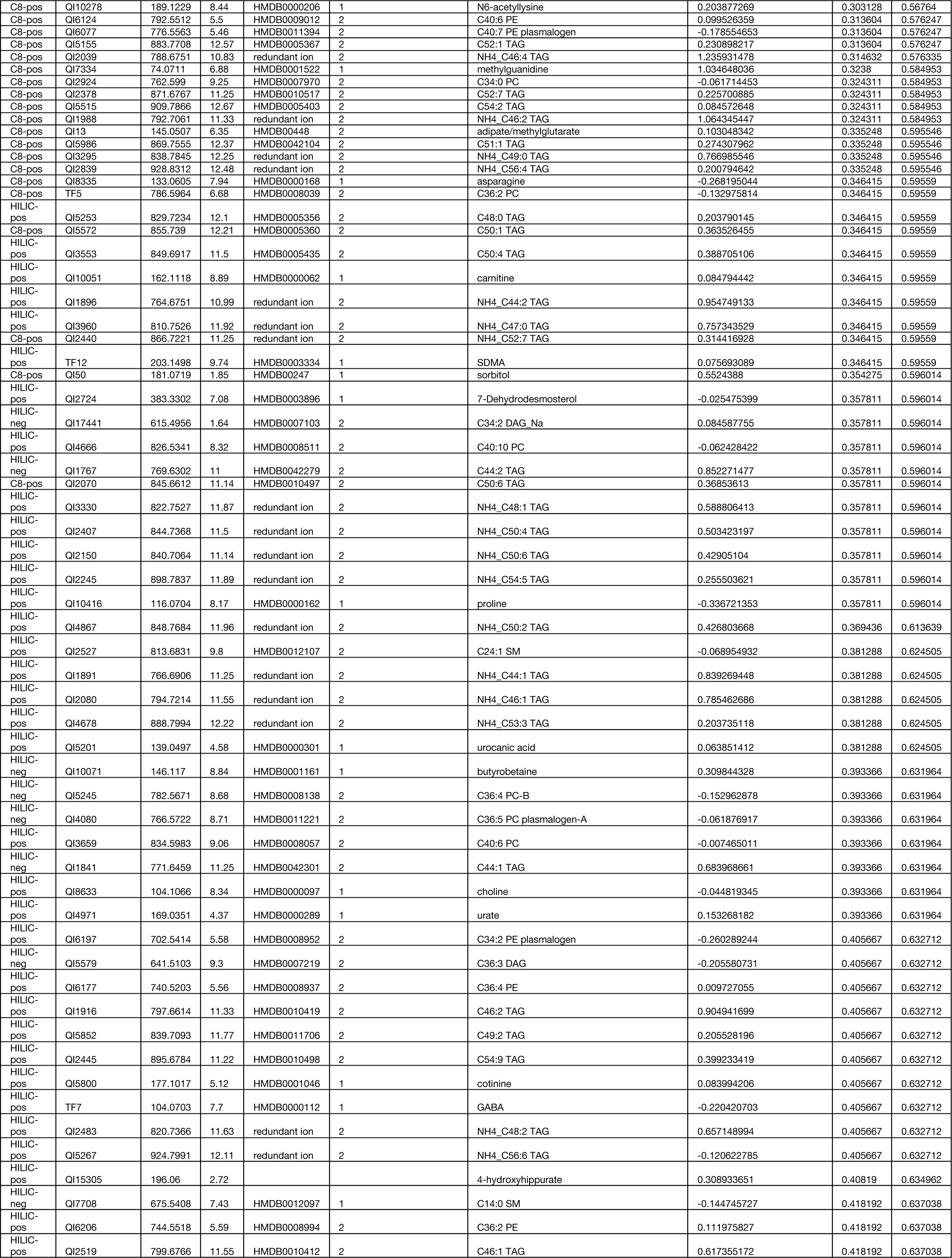
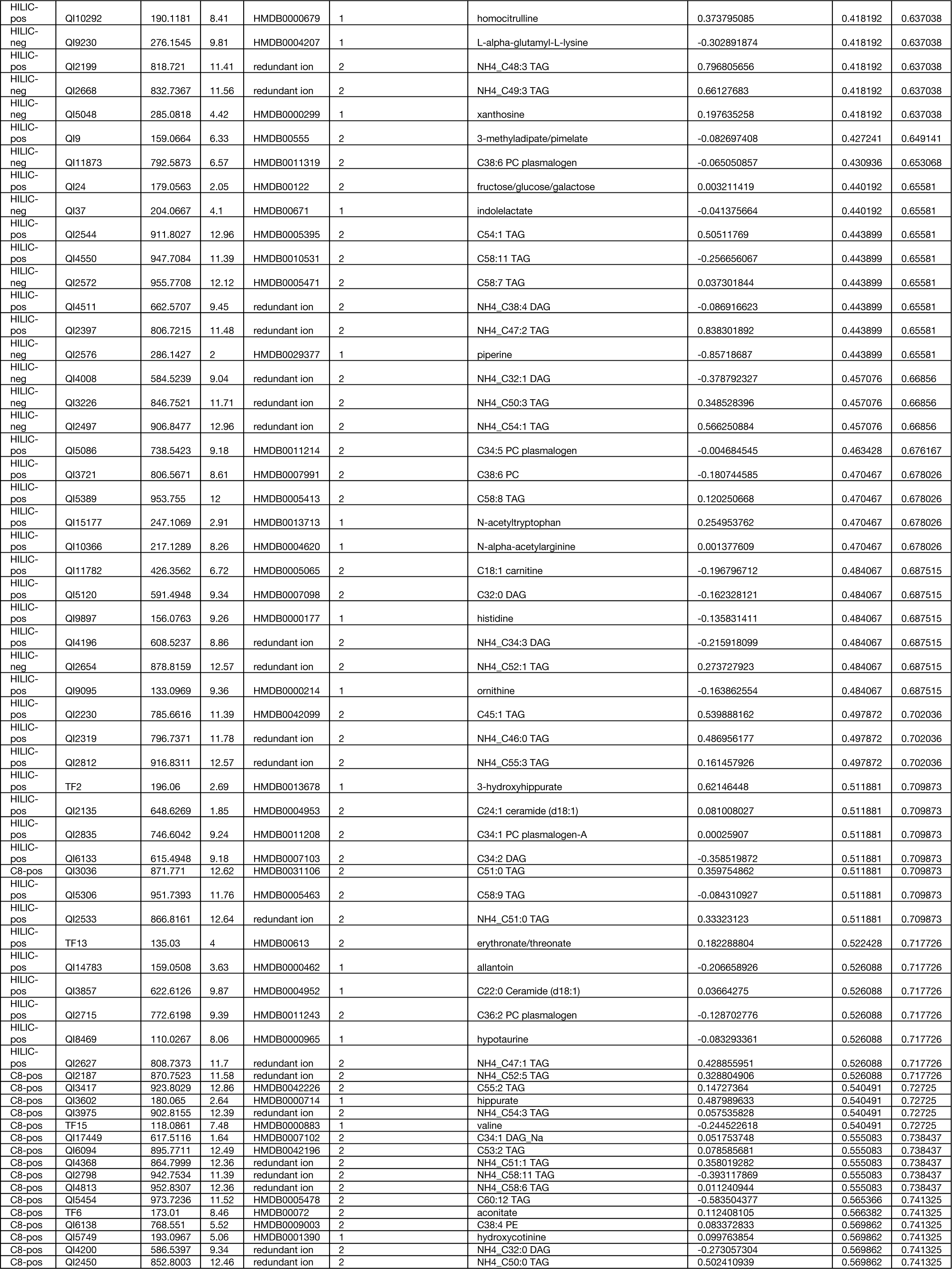
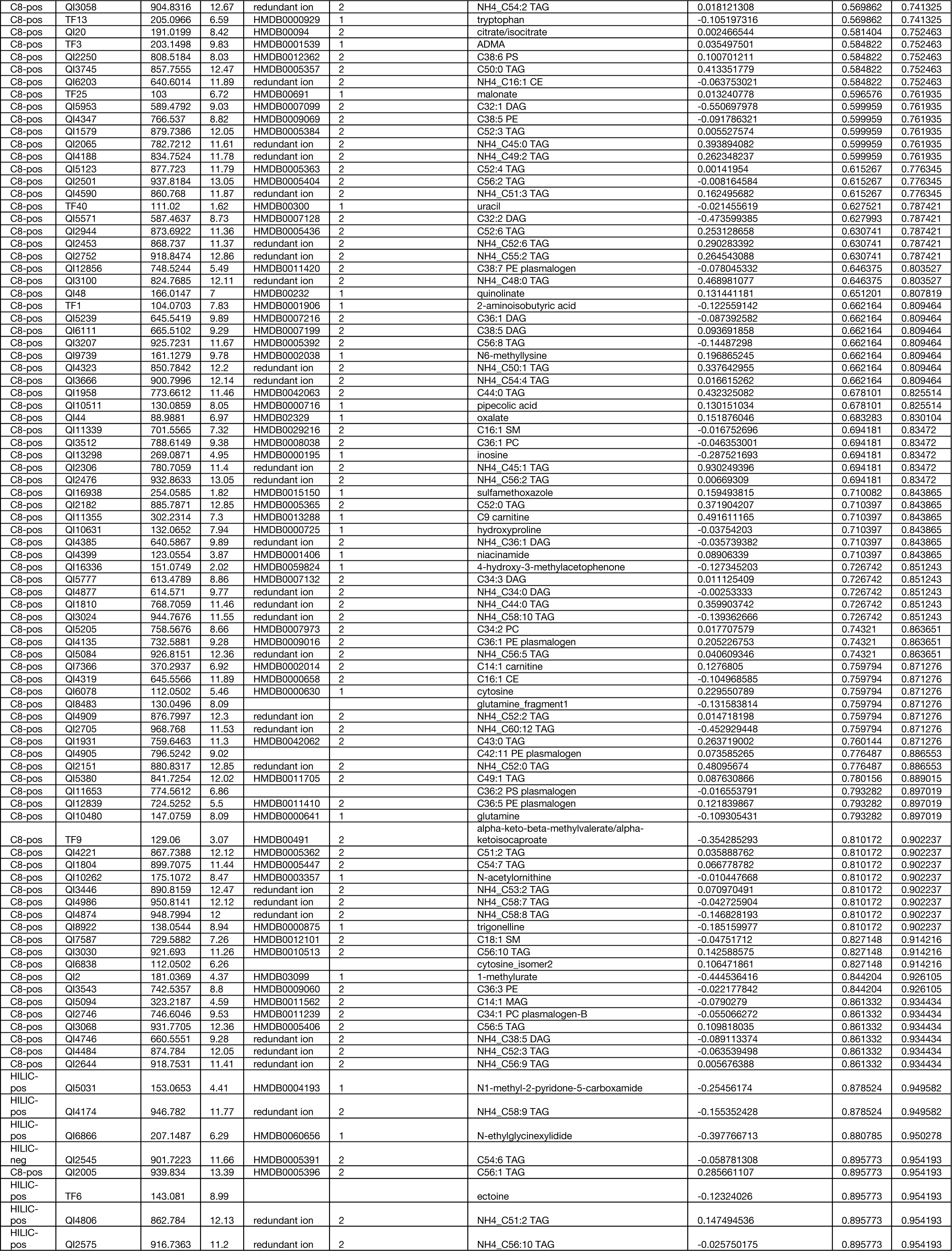
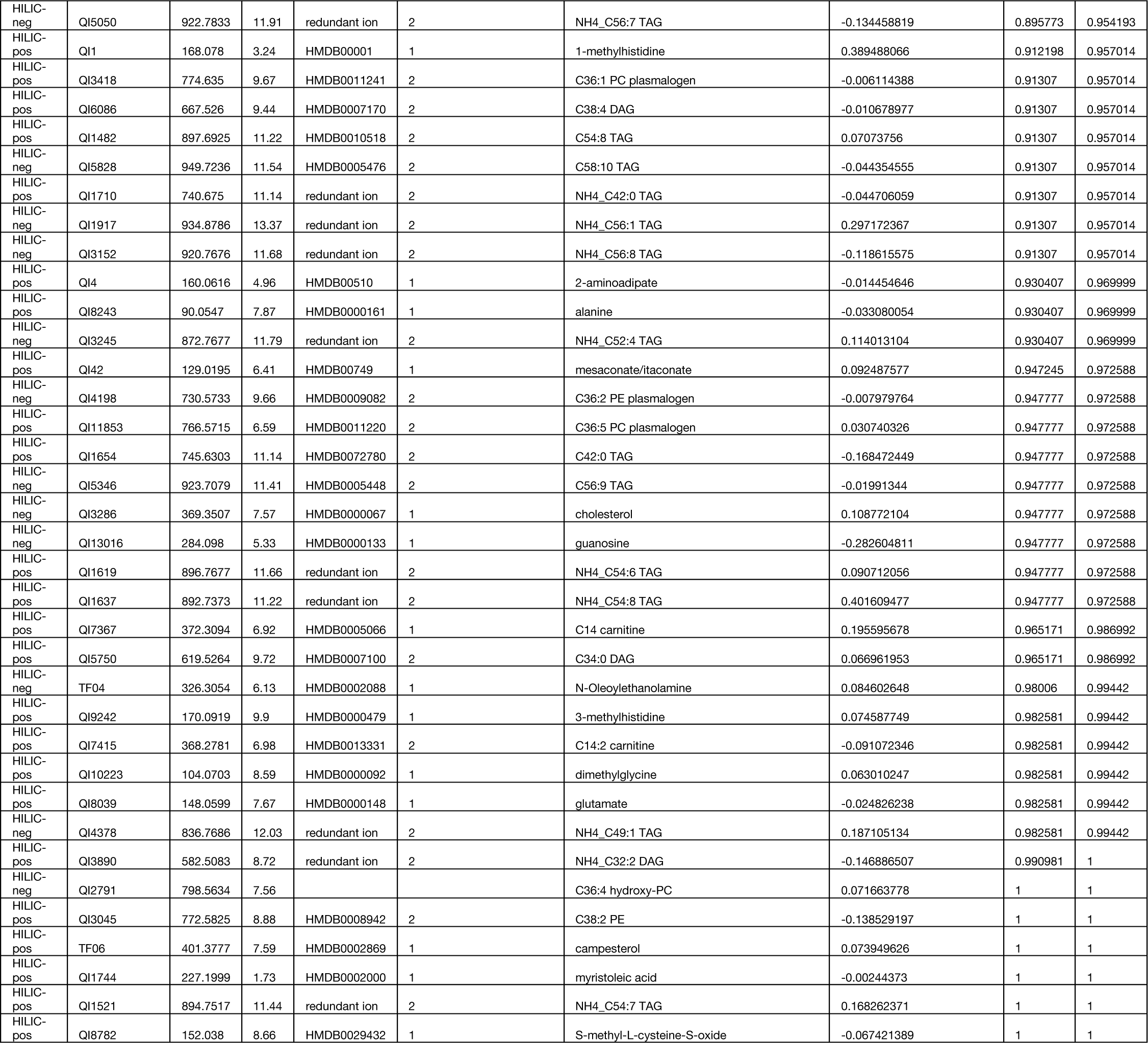
Targeted metabolomics results.

**Table S3.**
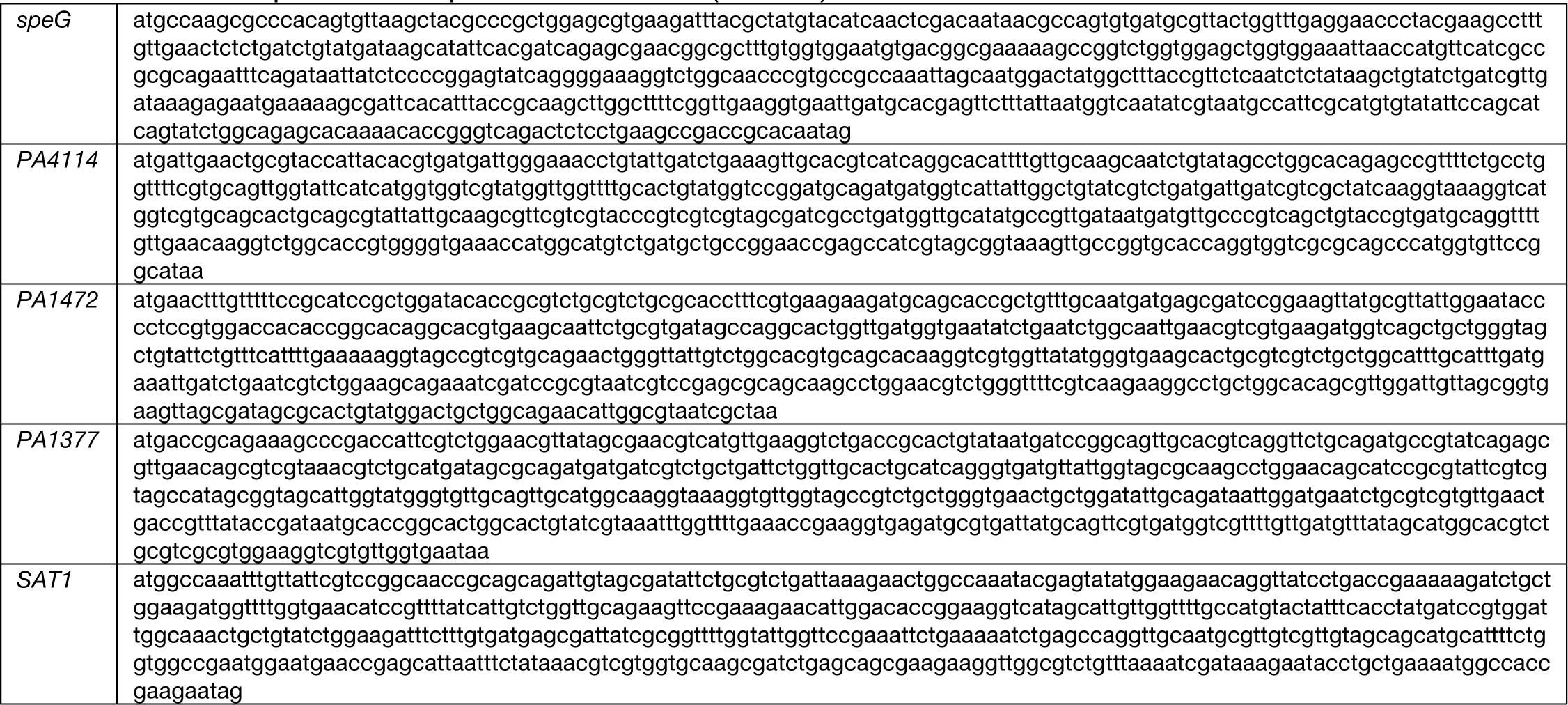
Codon optimized gene sequences for *E. coli* expression (5’ -> 3’)

**Table S4.**
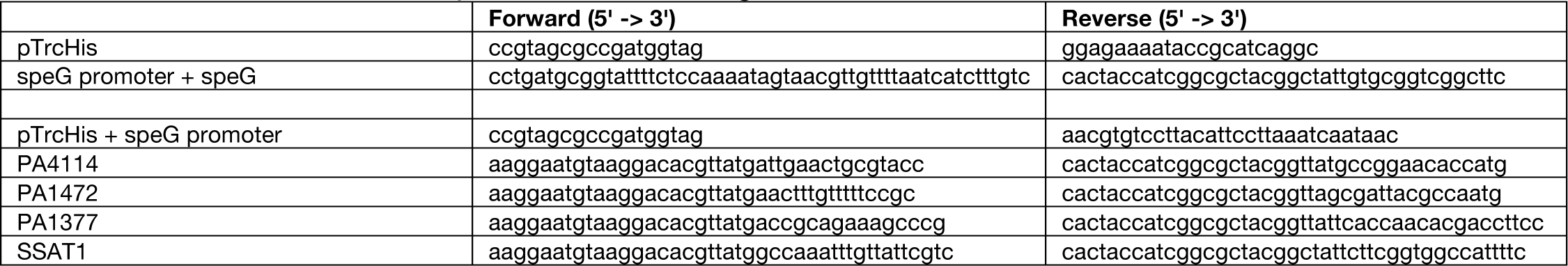
Primers used for complementation cloning.

**Table S5.**
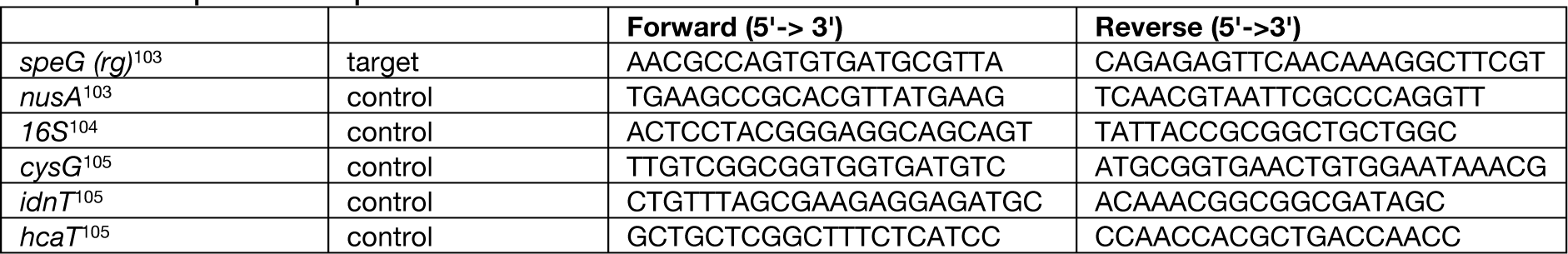
qRT-PCR primers.

**Table S6.**
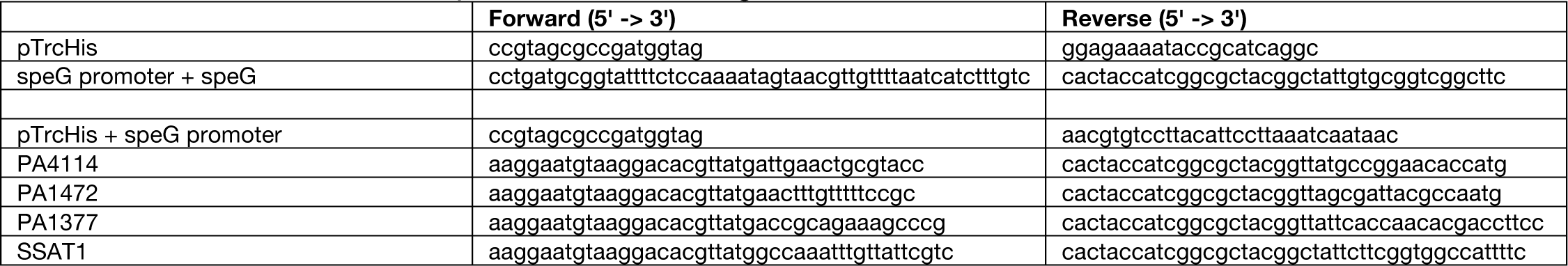
Primers used for complementation cloning.

**Table S7.**
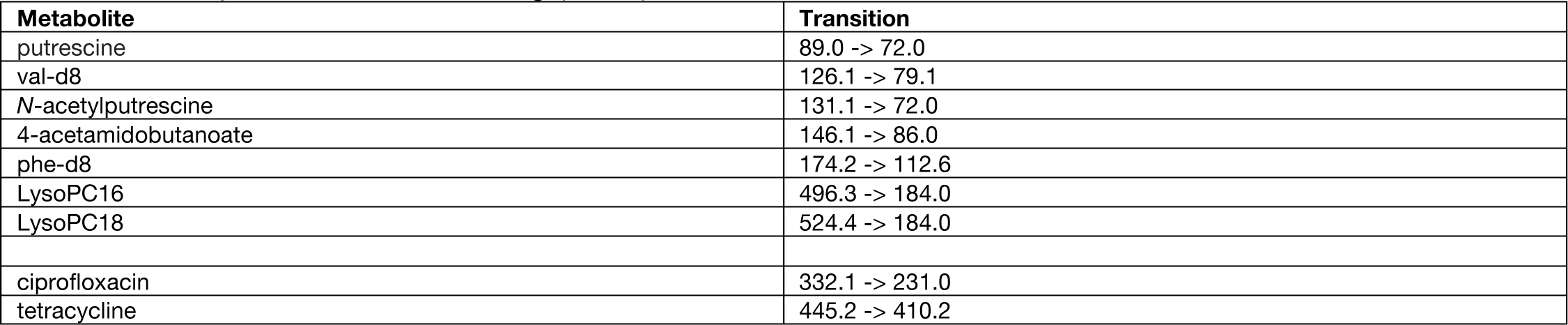
Multiple reaction monitoring (MRM) transitions.

| Metabolite | Transition |
| --- | --- |
| putrescine | 89.0 -> 72.0 |
| val-d8 | 126.1 -> 79.1 |
| *N*-acetylputrescine | 131.1 -> 72.0 |
| 4-acetamidobutanoate | 146.1 -> 86.0 |
| phe-d8 | 174.2 -> 112.6 |
| LysoPC16 | 496.3 -> 184.0 |
| LysoPC18 | 524.4 -> 184.0 |
| ciprofloxacin | 332.1 -> 231.0 |
| tetracycline | 445.2 -> 410.2 |

